# MMP-12 exacerbates neuroinflammation in the ischemic brain: possible interactions with tPA and progranulin

**DOI:** 10.64898/2026.11.23.768772

**Authors:** Siva Reddy Challa, Casimir A. Fornal, Isidra M. Baker, Pavani Unnam, Samantha N. Jackson, Vishesh Vinayagam, Joseph Nicestro, Vinay Sama, Aaron T. VanGeest, Kara R. Proctor, Khushi A. Raghunandan, Jerusha Boyineni, Jeffrey D. Klopfenstein, Raghu Vemuganti, Krishna Kumar Veeravalli

## Abstract

The molecular mechanisms underlying the protection mediated by matrix metalloproteinase-12 (MMP-12) suppression following cerebral ischemia and reperfusion remain unknown. We hypothesized that MMP-12 suppression attenuates neuroinflammation. Young male rats underwent transient middle cerebral artery occlusion (MCAO) followed by reperfusion. Stroke severity was determined using the modified neurological severity score. MMP-12 was knocked down by administering MMP-12 shRNA plasmids. MMP-12 exhibited the highest upregulation among all MMPs examined, and its expression positively correlated with stroke severity. MMP-12 expression also positively correlated with astrocyte A1-type and microglia/macrophage M1-type markers. MMP-12 suppression attenuated the expression of A1- and M1-type markers and altered the morphology of astrocytes and microglia/macrophages. Positive correlations were also observed between MMP-12 expression and markers of monocytes and neutrophils, and MMP-12 suppression significantly reduced the expression of myeloperoxidase. Additionally, our results suggested a possible interaction between MMP-12 and both tissue-type plasminogen activator (tPA) and progranulin (PGRN). MMP-12 suppression reduced tPA protein expression and altered PGRN protein expression patterns, suggesting changes in PGRN processing. In summary, reducing MMP-12 expression attenuated post-stroke neuroinflammation, possibly by reducing gliosis, astrocyte and microglia/macrophage activation, A1- and M1-type marker expression, and monocyte and neutrophil infiltration, as well as the expression of multiple MMPs and tPA and the processing of PGRN.

## Introduction

Following an acute ischemic stroke, which accounts for 87% of all strokes, recanalization treatments such as thrombolysis with tissue-type plasminogen activator (tPA) or tenecteplase and endovascular thrombectomy restore blood flow.^1^ While reperfusion is essential to salvage ischemic tissue, the sudden return of blood and oxygen can trigger reperfusion injury.

We previously reported the induction of several matrix metalloproteinases (MMPs) in the rodent brain following transient focal cerebral ischemia.^2^ MMP-12 exhibited the highest upregulation among all MMPs examined in the ischemic brain of both rats and mice.^2,3^ Several studies from other laboratories have also reported increased MMP-12 expression in the ischemic brain of mice subjected to transient focal ischemia.^4–6^ Furthermore, an MMP-12 gene polymorphism associated with increased MMP-12 expression in humans has been linked to an increased risk of ischemic stroke.^7^ In addition, a genome-wide association study showed that elevated serum MMP-12 levels in patients with diabetes were positively associated with stroke severity and poor neurological outcomes.^8^ A recent report highlights the emerging role of MMP-12 in secondary brain damage and impaired functional recovery in rodents following transient focal cerebral ischemia.^9^ MMP-12 suppression was shown to reduce infarct volume and blood-brain barrier (BBB) disruption and enhance the recovery of sensorimotor function following transient focal cerebral ischemia in rats.^2,10,11^ MMP-12 suppression also reduced post-stroke infarct volume, BBB disruption, and brain atrophy and improved the recovery of motor and cognitive functions in mice.^6^ However, the molecular mechanisms underlying the protection mediated by MMP-12 suppression remain unclear.

Given the reported involvement of MMP-12 in the degradation of extracellular matrix and BBB tight junction proteins, the processing of other MMPs and biomolecules such as pro-TNFα and progranulin (PGRN), and its potential interplay with endogenous tPA,^2,6,9,10,12^ we investigated the role of MMP-12 in neuroinflammation following transient focal cerebral ischemia.

## Methods

### Induction of transient focal cerebral ischemia

Male Sprague-Dawley rats (2‒3 months old; Envigo, Indianapolis, IN, USA) were subjected to transient 2-h middle cerebral artery occlusion (MCAO) under isoflurane anesthesia using the intraluminal monofilament suture technique, as described by our group.^13^ Rats were randomly assigned to the following experimental groups (Supplementary Table 1): Sham, Stroke (2-h MCAO), Ctrl-sh (stroke + control shRNA treatment), and M12-sh (stroke + MMP-12 shRNA treatment). Pre- and postsurgical care was provided in accordance with the Ischemia Models: Procedural Refinements of In Vivo Experiments (IMPROVE) guidelines.^14^ All animal procedures were approved by the Institutional Animal Care and Use Committee. All experiments were conducted and data were reported following the guidelines outlined in the *Guide for the Care and Use of Laboratory Animals* (NIH Publication No. 86-23, revised; U.S. Department of Health and Human Services) and Animal Research: Reporting of In Vivo Experiments (ARRIVE).^15^ Stroke severity was evaluated at 2‒4 h and on post-MCAO day 1 using the modified Neurological Severity Score (mNSS), as described earlier.^13^ Scores range from 0 (no deficit) to 18 (maximal neurological impairment). Rats exhibiting mNSS scores ≥ 8 at 2‒4 h or on post-MCAO day 1 were included in the experimental group comparisons. However, for correlation analyses, stroke animals with mNSS scores < 8 were also included to examine the relationships across a broader range of stroke severity.

### MMP-12 knockdown

Control shRNA and MMP-12 shRNA plasmids were designed and synthesized using the pSilencer™ 4.1-CMV neo vector as described earlier.^11^ Briefly, plasmids from bacterial cultures incubated overnight (∼16 h) were extracted using a plasmid maxi kit (Qiagen, Germantown, MD, USA). Nanoparticle formulations of shRNA plasmids were prepared using in vivo-jetPEI reagent (VWR, Radnor, PA, USA) and administered intravenously (1 mg/kg) via the tail vein within 30 min of reperfusion.

### Real-time PCR

A comprehensive description of the procedure is provided in the supplementary methods. Briefly, total RNA was extracted from the ipsilateral cerebral hemisphere collected on post-MCAO day 3 using TRIzol reagent (Thermo Fisher Scientific, Waltham, MA, USA). One microgram of total RNA from each sample was reverse transcribed into cDNA using the iScript cDNA Synthesis Kit (Bio-Rad Laboratories, Hercules, CA, USA). For each diluted (1:10) cDNA sample, reactions were assembled using iTaq Universal SYBR Green Supermix (Bio-Rad Laboratories, Hercules, CA, USA). Forward and reverse primers for the target genes (Integrated DNA Technologies, Coralville, IA, USA) were diluted 1:10 in nuclease-free water. Primer sequences are listed in Supplementary Table 2. Reactions were run on an iCycler iQ Multi-Color Real-Time PCR Detection System (Bio-Rad Laboratories, Hercules, CA, USA). Relative gene expression was normalized to 18S rRNA, and fold change relative to sham was calculated using ΔCt values.

### Immunoblotting

Protein samples (50 µg per lane) from the ipsilateral hemispheres collected on post-MCAO day 3 were electrophoresed, transferred to nitrocellulose membranes, and probed with primary antibodies against the target proteins (Supplementary Table 3), followed by the appropriate HRP-conjugated secondary antibodies. Immunoreactive bands were visualized using enhanced chemiluminescence (ECL) Western blot detection reagent (Thermo Fisher Scientific, Waltham, MA, USA) on a KwikQuant Digital Western Blot Detection System (Kindle Biosciences, Greenwich, CT, USA). Each membrane was stripped and reprobed with a mouse monoclonal anti-GAPDH antibody followed by an HRP-conjugated goat anti-mouse secondary antibody (Supplementary Table 3). Immunoreactive bands representing GAPDH were also visualized using the ECL reagent and the KwikQuant Detection System. Band intensities were quantified using NIH ImageJ software (version 1.54p) and normalized to GAPDH.

### Immunoprecipitation

MMP-12 was immunoprecipitated from the ipsilateral cerebral hemispheres collected on post-MCAO day 3 using the Dynabeads™ Protein A Immunoprecipitation Kit (Thermo Fisher Scientific, Waltham, MA, USA) and an anti-MMP-12 antibody (Supplementary Table 3). A comprehensive description of the procedure is provided in the supplementary methods. Both the collected flow-through fraction and the final eluate from each sample were subjected to immunoblot analysis, as described in the previous section, using a rabbit monoclonal anti-tPA antibody and an HRP-conjugated goat anti-rabbit secondary antibody (Supplementary Table 3).

### Immunofluorescence

A comprehensive description of the procedure is provided in the supplementary methods. Briefly, on post-MCAO day 3, rats were deeply anesthetized and perfused transcardially with 1× PBS followed by 10% neutral buffered formalin. Brains were extracted, postfixed, cryoprotected, and sectioned. The coronal brain sections were immunostained with primary antibodies followed by appropriate secondary antibodies (Supplementary Table 3) and counterstained with 4′,6-diamidino-2-phenylindole (DAPI; 1 µg/mL in 1× PBS). Images were acquired using a 40× objective on an Olympus FluoView FV3000 confocal laser scanning microscope. For each animal, one field of view within the peri-infarct region was imaged at a consistent anatomical location across animals. Identical imaging and channel settings were used for all samples to permit direct comparison of fluorescence staining among experimental groups. Quantification of immunofluorescence was performed using NIH ImageJ software (version 1.53e). Integrated density (arbitrary units) was used as the quantitative measure of immunofluorescence. Astrocyte and microglia/macrophage morphology and co-labeling of selected markers were also visually examined.

### Exclusion criteria

Rats that died or were euthanized when moribund, as well as those exhibiting hemorrhage in the MCA region upon postmortem examination, were excluded. Rats exhibiting mNSS scores < 8 at both 2‒4 h and on post-MCAO day 1 were also excluded from the experimental group comparisons. However, stroke animals with mNSS scores < 8 were retained for correlation analyses. Statistical outliers were identified using the ROUT test (Q = 0.1%) and excluded from the analysis, if present.

### Statistical analysis

Statistical analyses were performed using GraphPad Prism 10.4.2 for Windows (GraphPad Software, San Diego, CA, USA). A comprehensive description of the statistical analyses and tests applied is provided in the supplementary methods. Quantitative data were tested for normality and homogeneity of variance. Based on the outcomes of these tests, appropriate statistical tests were applied. Differences between groups were considered significant at *p* < 0.05 or false discovery rate (FDR)-adjusted *p* < 0.05, as appropriate. Supplementary Table 4 summarizes the statistical analyses, including the tests applied, test statistics (t ratios and t- and F-values), and exact *p*-values.

## Results

### MMP-12 expression increases in the ischemic brain

Following transient focal cerebral ischemia, mRNA expression of all 22 MMPs examined was increased in the ipsilateral hemisphere (Fig. 1A). Eleven MMPs exhibited ≥10-fold increases, and among these, MMP-12 exhibited the greatest upregulation (124-fold relative to Sham). MMP-12 mRNA expression was positively correlated with mNSS (Fig. 1B). MMP-12 protein expression was also significantly increased in the ipsilateral hemisphere compared with Sham (Fig. 1C and D). Rats in the M12-sh group exhibited significantly reduced MMP-12 protein expression compared with those in the Ctrl-sh group following transient focal cerebral ischemia (Fig. 1E).

**Fig. 1.**
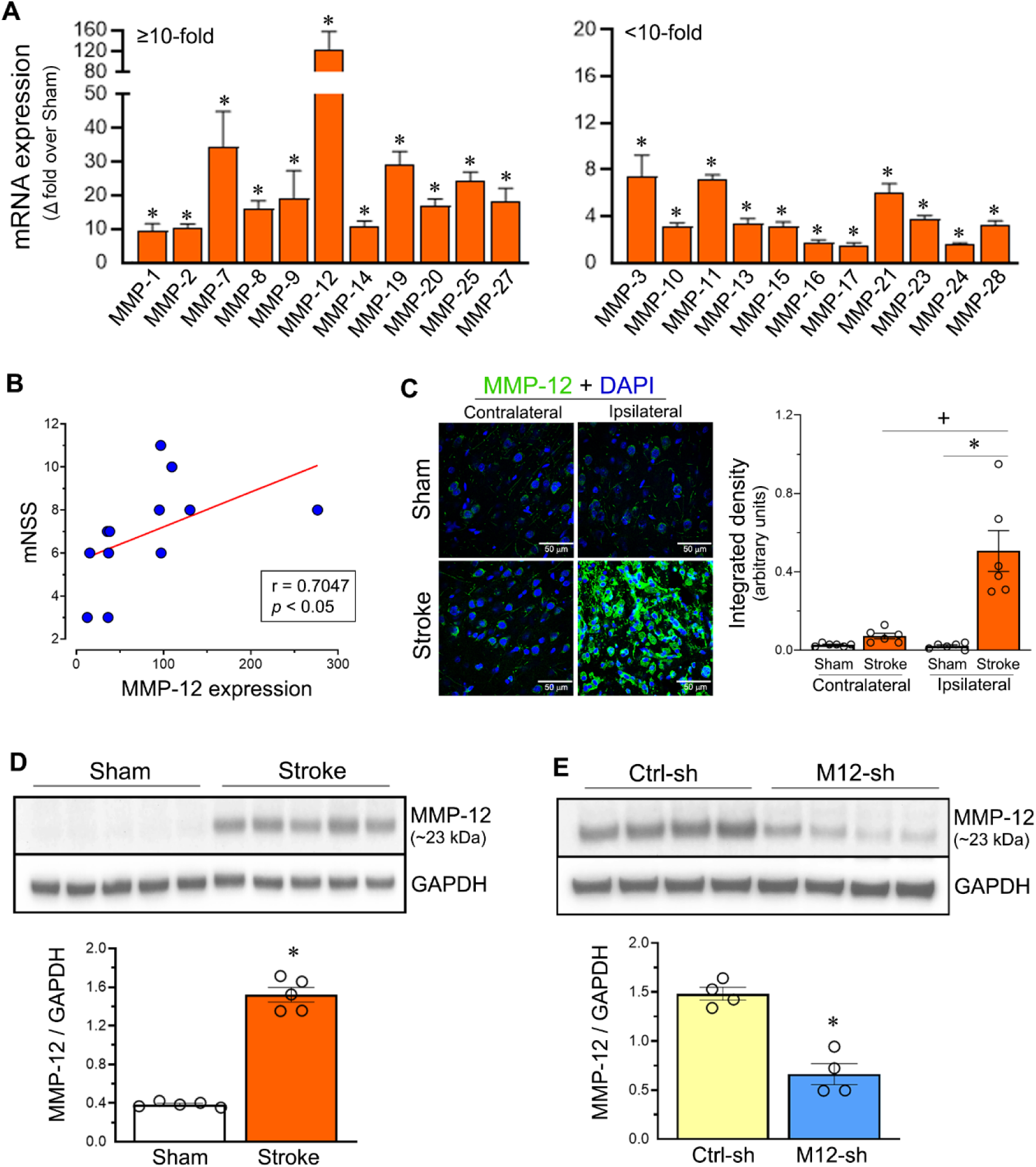
MMP-12 expression increases in the brain following cerebral I/R. (**A**) Bar graph shows the relative mRNA expression of 22 MMPs in the ipsilateral hemisphere of rats on post-MCAO day 3. Values above each bar indicate the fold change. Data are presented as mean ± SEM; n = 6 per group. \**p* < 0.05 vs. Sham group. (**B**) Scatter plot shows a significant positive Spearman correlation between the modified neurological severity score (mNSS) assessed 2‒4 h after MCAO and relative MMP-12 mRNA expression in the ipsilateral hemisphere on post-MCAO day 3. n = 12. (**C**) Representative images showing MMP-12 protein expression (green) in the contralateral and ipsilateral hemispheres of rats from Sham and Stroke groups on post-MCAO day 3. In the Stroke group, images of the ipsilateral hemisphere were obtained from the peri-infarct cortex. Nuclei (blue) were stained with DAPI. Column scatter plot shows the integrated density of green fluorescence corresponding to MMP-12 protein expression in the contralateral and ipsilateral hemispheres of rats from the Sham and Stroke groups. n = 6 per group. \**p* < 0.05 and ^+^*p* < 0.05. (**D, E**) Immunoblots show the protein expression of MMP-12 and GAPDH in the ipsilateral hemisphere of rats from the Sham, Stroke, Ctrl-sh (stroke + control shRNA treatment), and M12-sh (stroke + MMP-12 shRNA treatment) groups on post-MCAO day 3. Column scatter plots show MMP-12 protein expression normalized to GAPDH. n = 4–5 per group. \**p* < 0.05 vs. Sham group (**D**). \**p* < 0.05 vs. Ctrl-sh group (**E**).

### MMP-12 suppression attenuates astrogliosis and reduces A1-type marker expression

Following transient focal cerebral ischemia, glial fibrillary acidic protein (GFAP) immunostaining was significantly increased in the peri-infarct cortex compared with Sham (Fig. 2A). In addition, pronounced astrocyte enlargement was observed in the Stroke group, consistent with astrocyte activation and astrogliosis. MMP-12 knockdown reduced astrocyte enlargement and significantly decreased GFAP protein expression in the ischemic brain (Fig. 2A).

**Fig. 2.**
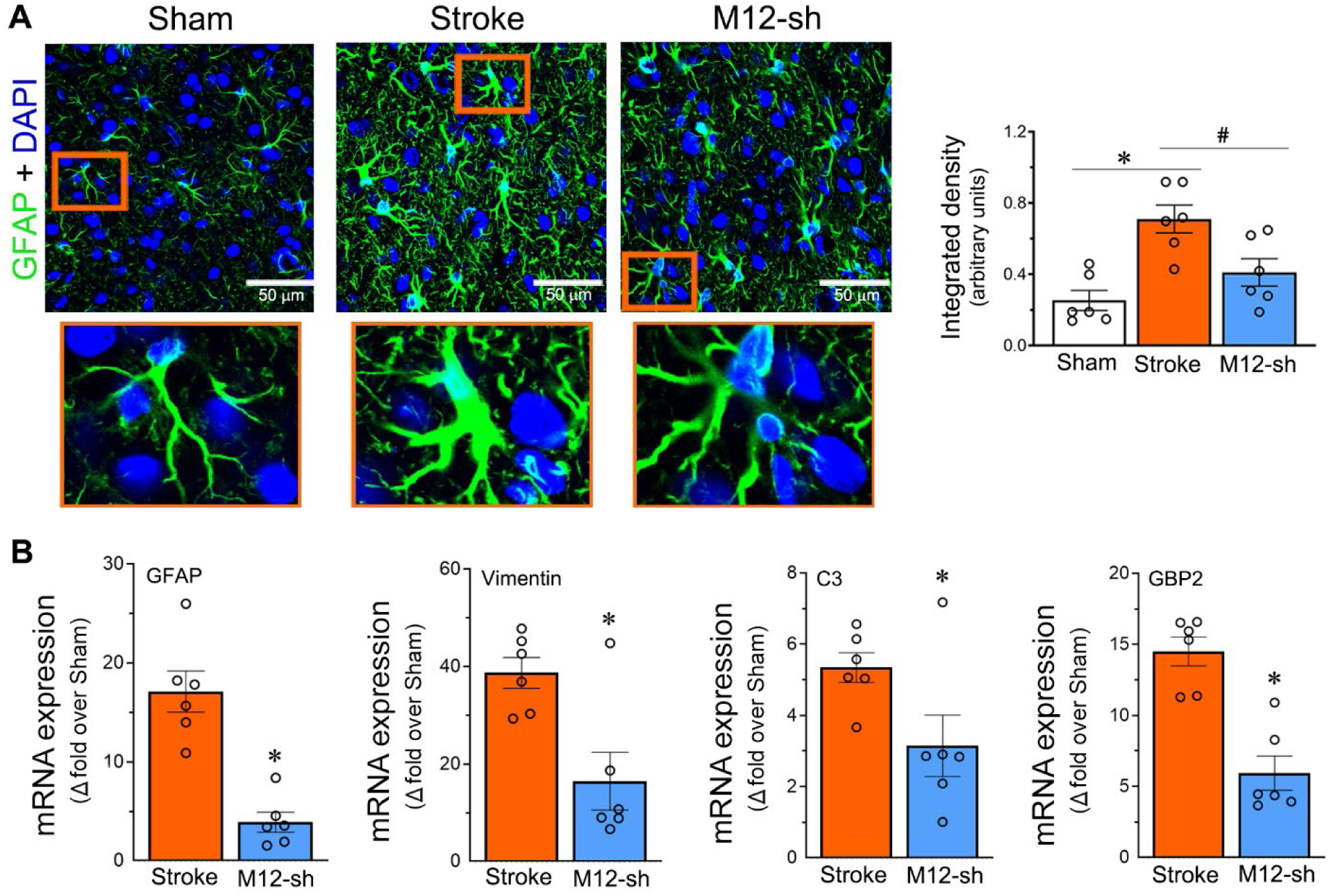
MMP-12 suppression following cerebral I/R alters astrocyte morphology and reduces astrocyte marker expression in the ischemic brain. (**A**) Representative images showing GFAP protein expression (green) in the peri-infarct cortex of the ipsilateral hemisphere of rats from the Sham, Stroke, and M12-sh (stroke + MMP-12 shRNA treatment) groups on post-MCAO day 3. The images in the bottom row are enlarged views of the boxed regions in the upper row. Nuclei (blue) were stained with DAPI. Column scatter plot shows the integrated density of green fluorescence corresponding to GFAP protein expression. Data are presented as mean ± SEM; n = 6 per group.\**p* < 0.05 and ^#^*p* < 0.05. (**B**) Column scatter plots show the relative mRNA expression of the astrocyte-associated markers GFAP and vimentin and the proinflammatory A1-type markers C3 and GBP2 in the ipsilateral hemisphere of rats from the Stroke and M12-sh groups on post-MCAO day 3. n = 6 per group. \**p* < 0.05 vs. Stroke group.

The mRNA expression of the astrocyte-associated markers GFAP and vimentin, as well as the proinflammatory A1-type markers complement component 3 (C3), guanylate-binding protein 2 (GBP2), and FK506 binding protein 5 (FKBP5), was significantly increased in the ipsilateral hemisphere (Supplementary Fig. 1). The mRNA expression levels of GFAP, vimentin, C3, and GBP2 were positively correlated with MMP-12 mRNA expression (Supplementary Fig. 2). MMP-12 knockdown significantly reduced the mRNA expression of these astrocyte-associated and proinflammatory A1-type markers in the ischemic brain (Fig. 2B).

### MMP-12 suppression reduces microglia/macrophage M1- and M2-type marker expression

Following transient focal cerebral ischemia, immunostaining for the microglia/macrophage marker ionized calcium-binding adaptor molecule 1 (Iba1) revealed activated microglia/macrophages with short, thick processes and enlarged cell bodies in the Stroke group, in contrast to the long, thin processes and relatively small cell bodies observed in the Sham group (Fig. 3A). MMP-12 knockdown did not reduce Iba1 expression compared with the Stroke group (Fig. 3B). However, microglia/macrophages in the M12-sh group exhibited an intermediate morphology, with moderately thickened processes and intermediate-sized cell bodies compared with those in the Sham and Stroke groups (Fig. 3A). These findings suggest that MMP-12 shRNA treatment may have attenuated microglia/macrophage activation.

**Fig. 3.**
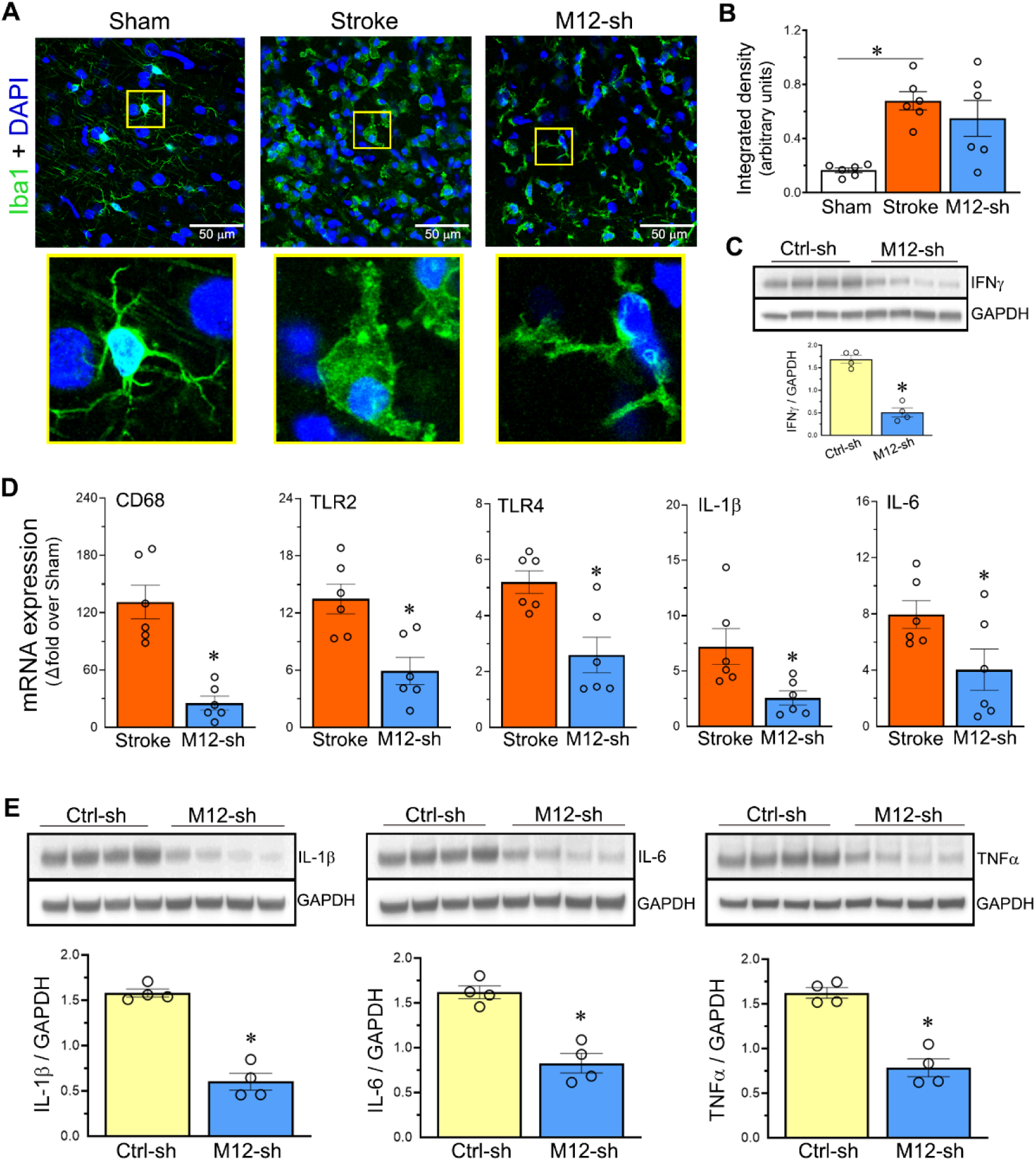
MMP-12 suppression following cerebral I/R alters microglia/macrophage morphology and reduces the expression of M1-type markers in the ischemic brain. (**A**) Representative images showing Iba1 protein expression (green) in the per-infarct region of the ipsilateral hemisphere of rats from the Sham, Stroke, and M12-sh (stroke + MMP-12 shRNA treatment) groups on post-MCAO day 3. The images in the bottom row are enlarged views of the boxed regions in the upper row. Nuclei (blue) were stained with DAPI. n = 6 per group. (**B**) Column scatter plot shows the integrated density of green fluorescence corresponding to Iba1 protein expression. Data are presented as mean ± SEM; n = 6 per group. \**p* < 0.05. ns, not significant. (**C**) Immunoblots show the protein expression of IFNγ and GAPDH in the ipsilateral hemisphere of rats from the Ctrl-sh (stroke + control shRNA treatment) and M12-sh groups on post-MCAO day 3. Column scatter plot shows IFNγ protein expression normalized to GAPDH. n = 4 per group. \**p* < 0.05 vs. Ctrl-sh group. (**D**) Column scatter plots show the relative mRNA expression of the M1-type proinflammatory markers CD68, TLR2, TLR4, IL-1β, and IL-6 in the ipsilateral and contralateral hemispheres of rats from the Stroke and M12-sh groups on post-MCAO day 3. n = 6 per group. \**p* < 0.05 vs. Stroke group. ns, not significant. (**E**) Immunoblots show the protein expression of the proinflammatory cytokines IL-1β, IL-6, and TNFα and GAPDH in the ipsilateral hemisphere of rats from the Ctrl-sh and M12-sh groups on post-MCAO day 3. Column scatter plots show IL-1β, IL-6, and TNFα protein expression normalized to GAPDH. n = 4 per group. \**p* < 0.05 vs. Ctrl-sh group.

IFNγ protein expression increased substantially in the Stroke group compared with the Sham group, consistent with increased proinflammatory M1-type activation (Supplementary Fig. 3). Similarly, mRNA expression of the M1-type proinflammatory markers CD68, CD16, Toll-like receptor 2 (TLR2), TLR4, inducible nitric oxide synthase (iNOS), C-C motif chemokine ligand 2 (CCL2), also known as monocyte chemoattractant protein-1 (MCP-1), C-X-C motif chemokine ligand 1 (CXCL1), interleukin-1β (IL-1β), IL-6, and tumor necrosis factor α (TNFα), was significantly increased in the ipsilateral brain following transient focal cerebral ischemia (Supplementary Fig. 4). Protein expression of the M1-type proinflammatory cytokines IL-1β, IL-6, and TNFα was also significantly increased in the ipsilateral brain of rats in the Stroke group compared with the Sham group (Supplementary Fig. 5). MMP-12 mRNA expression was positively correlated with CCL2, CXCL1, and IL-6 mRNA expression, demonstrating an association between increased MMP-12 expression and increased proinflammatory M1-type marker expression (Supplementary Fig. 6). IFNγ protein expression, which is associated with the induction of proinflammatory M1-type microglia/macrophage activation, was significantly reduced by MMP-12 shRNA treatment compared with control shRNA treatment (Fig. 3C). As expected, MMP-12 shRNA treatment also decreased mRNA expression of the M1-type markers CD68, TLR2, TLR4, IL-1β, and IL-6 compared with the Stroke group and protein expression of the M1-type proinflammatory cytokines IL-1β, IL-6, and TNFα compared with control shRNA treatment (Fig. 3D and E).

The mRNA expression levels of the M2-type anti-inflammatory markers CD163, CD206, arginase 1 (Arg1), IL-10, and transforming growth factor β (TGFβ) were significantly upregulated in the Stroke group compared with the Sham group (Supplementary Fig. 7) and were positively correlated with MMP-12 mRNA expression (Supplementary Fig. 8). Contrary to our expectation, MMP-12 knockdown significantly decreased, rather than increased, the mRNA expression of the M2-type markers CD163, CD206, Arg1, IL-10, and TGFβ compared with the Stroke group (Supplementary Fig. 9).

### MMP-12 expression is associated with monocyte and neutrophil markers

Transient focal cerebral ischemia significantly increased the mRNA expression levels of C-C motif chemokine receptor 2 (CCR2; expressed by monocytes and macrophages), C-X-C Motif chemokine receptor 2 (CXCR2; expressed by neutrophils), elastase, neutrophil expressed (ELANE), and myeloperoxidase (MPO; expressed mainly by neutrophils and, to a lesser extent, monocytes) (Fig. 4A). MMP-12 mRNA expression was positively correlated with CCR2, CXCR2, and MPO mRNA expression (Fig. 4B). MPO protein expression was also significantly increased in the Stroke group compared with the Sham group (Fig. 4C). MMP-12 knockdown significantly reduced MPO protein expression compared with control shRNA treatment (Fig. 4D).

**Fig. 4.**
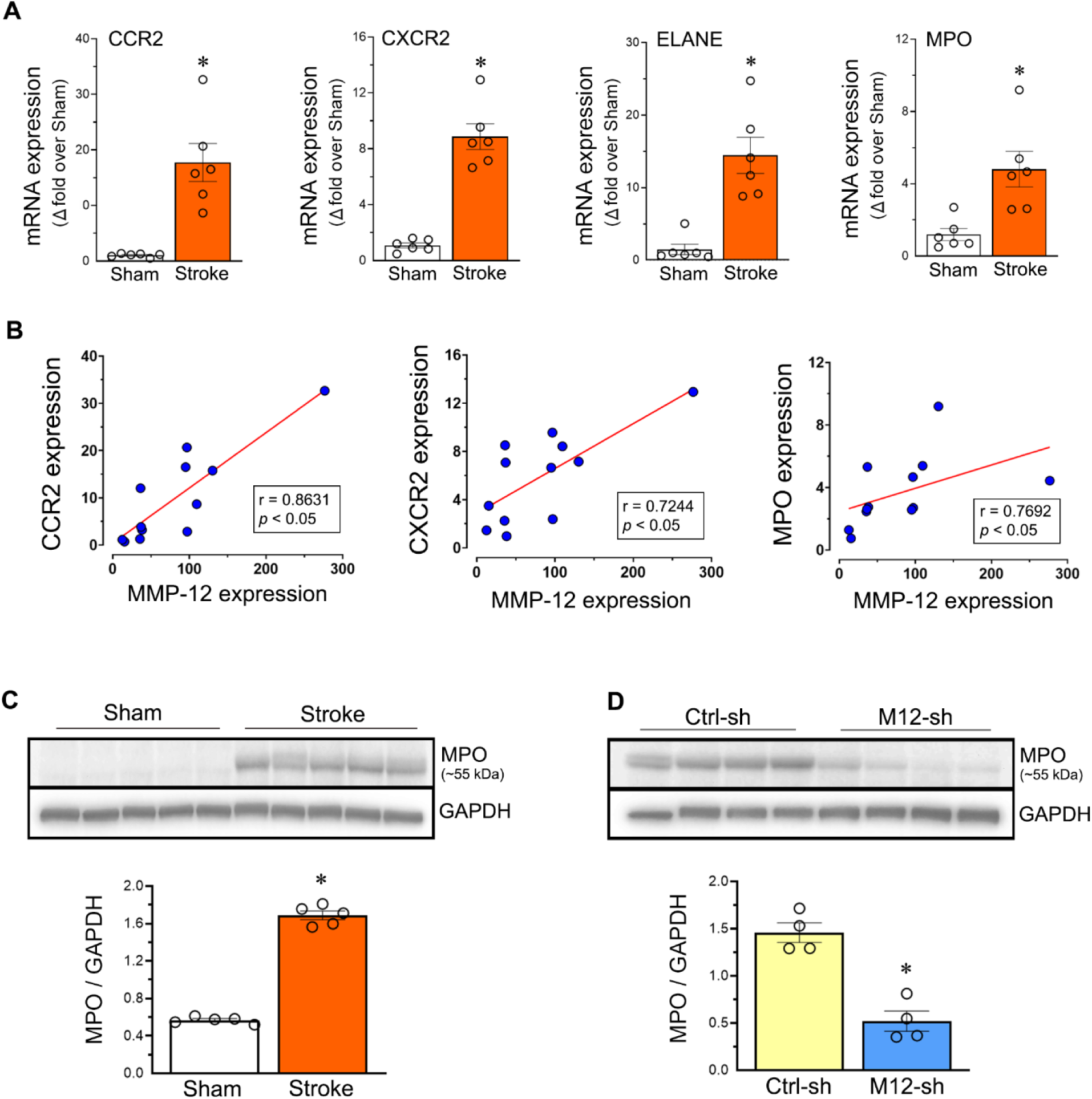
MMP-12 expression in the ischemic brain is associated with monocyte/macrophage- and neutrophil-associated markers. (**A**) Column scatter plots show the relative mRNA expression of the monocyte/macrophage- and neutrophil-associated markers CCR2, CXCR2, ELANE, and MPO in the ipsilateral hemisphere of rats from the Sham and Stroke groups on post-MCAO day 3. Data are presented as mean ± SEM; n = 6 per group. \**p* < 0.05 vs. Sham group. (**B**) Scatter plots show positive Pearson or Spearman correlations, as appropriate, between relative MMP-12 mRNA expression and relative mRNA expression of CCR2, CXCR2, and MPO in the ipsilateral hemisphere of rats on post-MCAO day 3. n = 12. (**C**) Immunoblots show the protein expression of MPO and GAPDH in the ipsilateral hemisphere of rats from the Sham and Stroke groups on post-MCAO day 3. Column scatter plot shows MPO protein expression normalized to GAPDH. n = 5 per group. \**p* < 0.05 vs. Sham group. (**D**) Immunoblots show the protein expression of MPO and GAPDH in the ipsilateral hemisphere of rats from the Ctrl-sh (stroke + control shRNA treatment) and M12-sh (stroke + MMP-12 shRNA treatment) groups on post-MCAO day 3. Column scatter plot shows MPO protein expression normalized to GAPDH. n = 4 per group. \**p* < 0.05 vs. Ctrl-sh group.

### MMP-12 suppression reduces the expression of other MMPs in the ischemic brain

Rats in the M12-sh group exhibited significantly reduced mRNA expression of MMP-2, MMP-3, MMP-7, MMP-8, MMP-9, MMP-11, MMP-14, MMP-15, MMP-19, MMP-21, MMP-23, MMP-25, MMP-27, and MMP-28, which were upregulated ≥ 2-fold in the ischemic brain, compared with the Stroke group (Fig. 5). These findings suggest that the expression of these MMPs may be regulated downstream of MMP-12 in the ischemic brain.

**Fig. 5.**
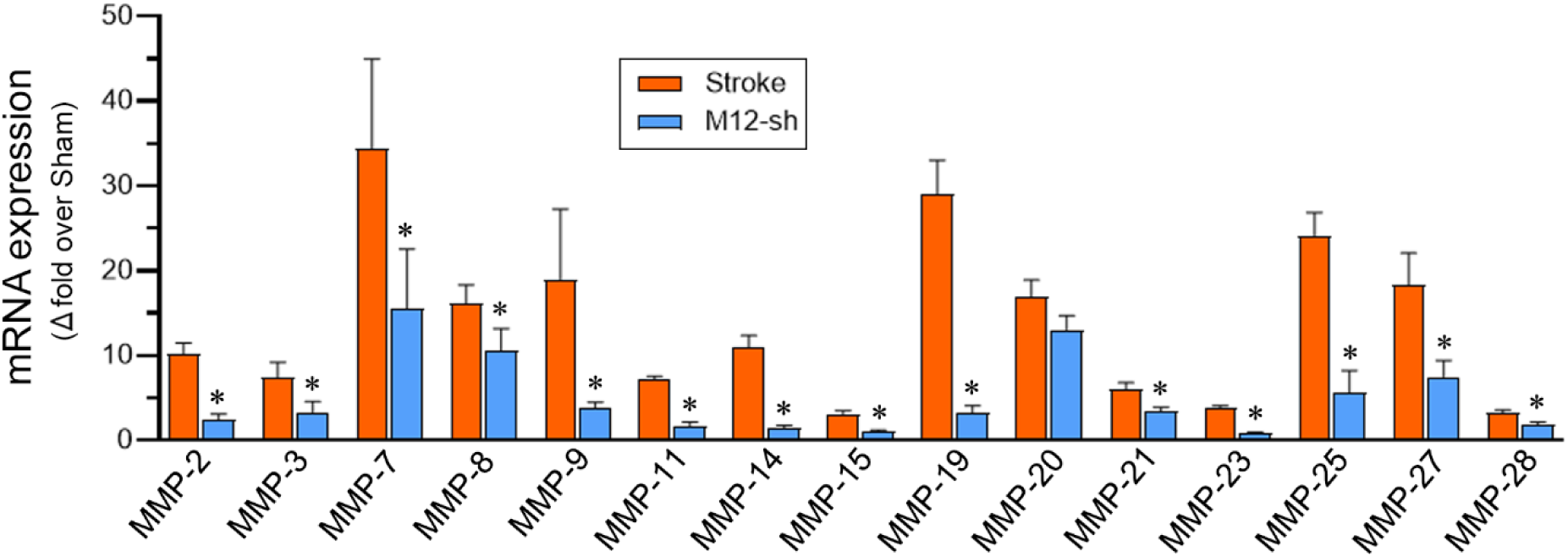
MMP-12 suppression reduces the expression of other MMPs in the ischemic brain. Bar graph shows the relative mRNA expression of 15 MMPs other than MMP-12 in the ipsilateral hemisphere of rats from the Stroke and M12-sh groups on post-MCAO day 3. Data are presented as mean ± SEM; n = 6 per group. \**p* < 0.05 vs. Stroke group.

### MMP-12 and tPA colocalize in the peri-infarct cortex

We recently reported the interplay between MMP-12 and tissue-type plasminogen activator (tPA) in the brain following transient focal cerebral ischemia.^12^ In the current study, MMP-12 and tPA exhibited prominent colocalization in the peri-infarct cortex of rats subjected to transient focal cerebral ischemia compared with the Sham group (Fig. 6A). Furthermore, tPA was co-immunoprecipitated with MMP-12 from ipsilateral brain tissue samples (Fig. 6B), supporting a possible molecular interaction between MMP-12 and tPA in the brain following transient focal cerebral ischemia. We previously reported that suppression of ischemia-induced endogenous tPA expression significantly attenuated neuroinflammation and BBB disruption.^12^ In the current study, endogenous tPA protein expression was significantly reduced in the M12-sh group rats compared with those in the Ctrl-sh group following transient focal cerebral ischemia (Fig. 6C). These findings suggest that MMP-12 may contribute to neuroinflammation in the ischemic brain, at least in part, through its association with endogenous tPA.

**Fig. 6.**
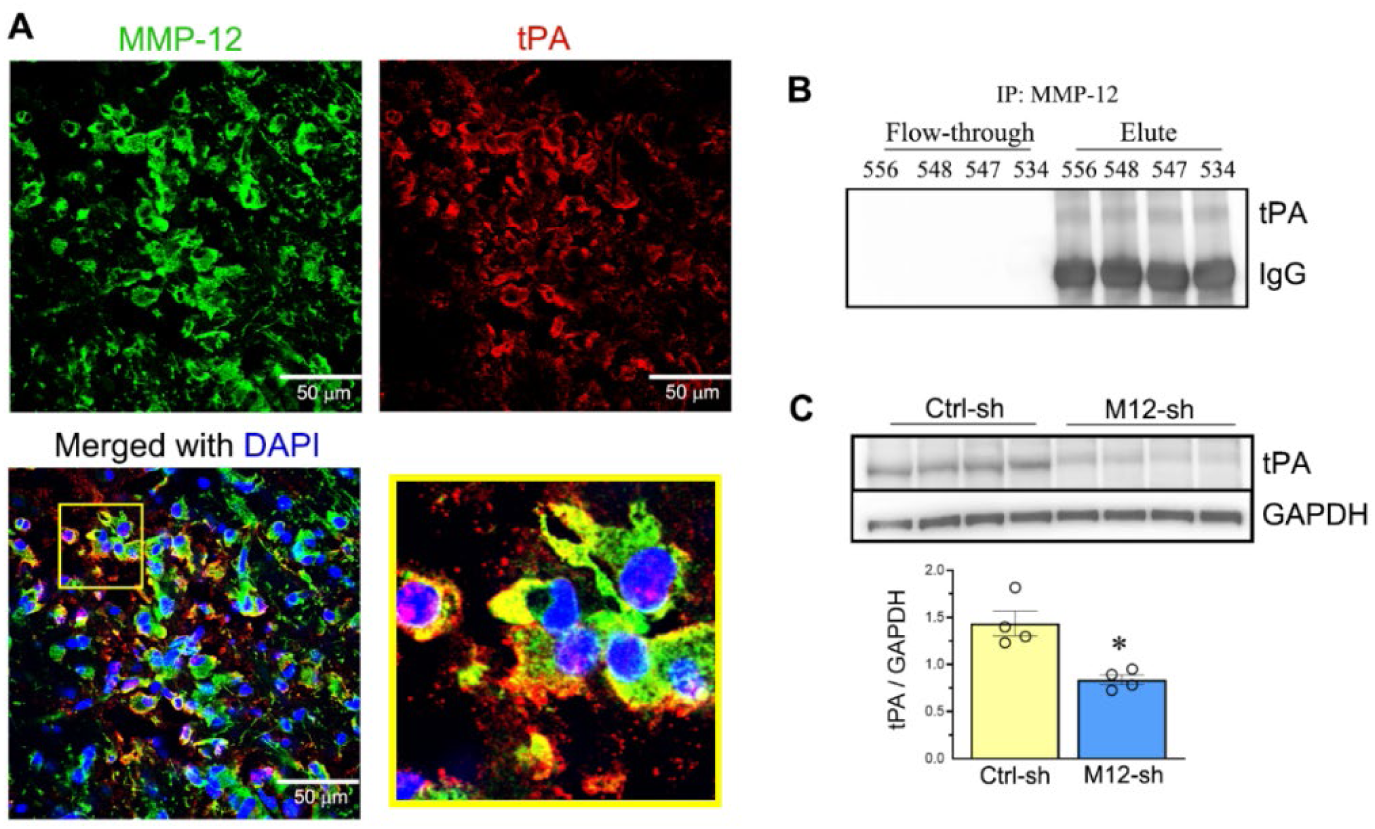
MMP-12 and tPA colocalize in the ischemic brain, and MMP-12 suppression reduces endogenous tPA expression. (**A**) Representative double-labeled immunofluorescence images showing MMP-12 protein expression (green), tPA protein expression (red), and their colocalization in the peri-infarct cortex of the ipsilateral hemisphere of rats from the Stroke group on post-MCAO day 3. Nuclei (blue) were stained with DAPI. n = 5. (**B**) Immunoprecipitation (IP) with an anti-MMP-12 antibody was used to immunoprecipitate MMP-12 from the ipsilateral hemisphere of rats from the Stroke group on post-MCAO day 3. Immunoblotting of the flow-through and eluate fractions obtained following immunoprecipitation detected tPA along with IgG in the eluate fraction, providing evidence of an association between MMP-12 and tPA in the ischemic brain. n = 4. (**C**) Immunoblots show the protein expression of tPA and GAPDH in the ipsilateral hemisphere of rats from the Ctrl-sh (stroke + control shRNA treatment) and M12-sh (stroke + MMP-12 shRNA treatment) groups on post-MCAO day 3. Column scatter plot shows tPA protein expression normalized to GAPDH. Data are presented as mean ± SEM; n = 4 per group. \**p* < 0.05 vs. Ctrl-sh group.

### MMP-12 suppression alters PGRN expression in the ischemic brain

Progranulin (PGRN) is highly expressed in microglia and neurons.^16–18^ PGRN is typically secreted as a highly glycosylated protein. PGRN mRNA expression increased significantly following transient focal cerebral ischemia compared with the Sham group (Fig. 7A). Two protein bands, at ∼80 and ∼68 kDa, were observed for PGRN (Fig. 7B). The ∼80-kDa band may represent a mature, glycosylated form of PGRN associated with neuroprotective and anti-inflammatory properties, whereas the ∼68-kDa band may represent a less extensively glycosylated or unmodified form of PGRN.^19^ Protein levels of the ∼68-kDa PGRN band, but not the ∼80-kDa band, increased significantly in the ischemic brain following transient focal cerebral ischemia compared with the Sham group (Fig. 7B).

**Fig. 7.**
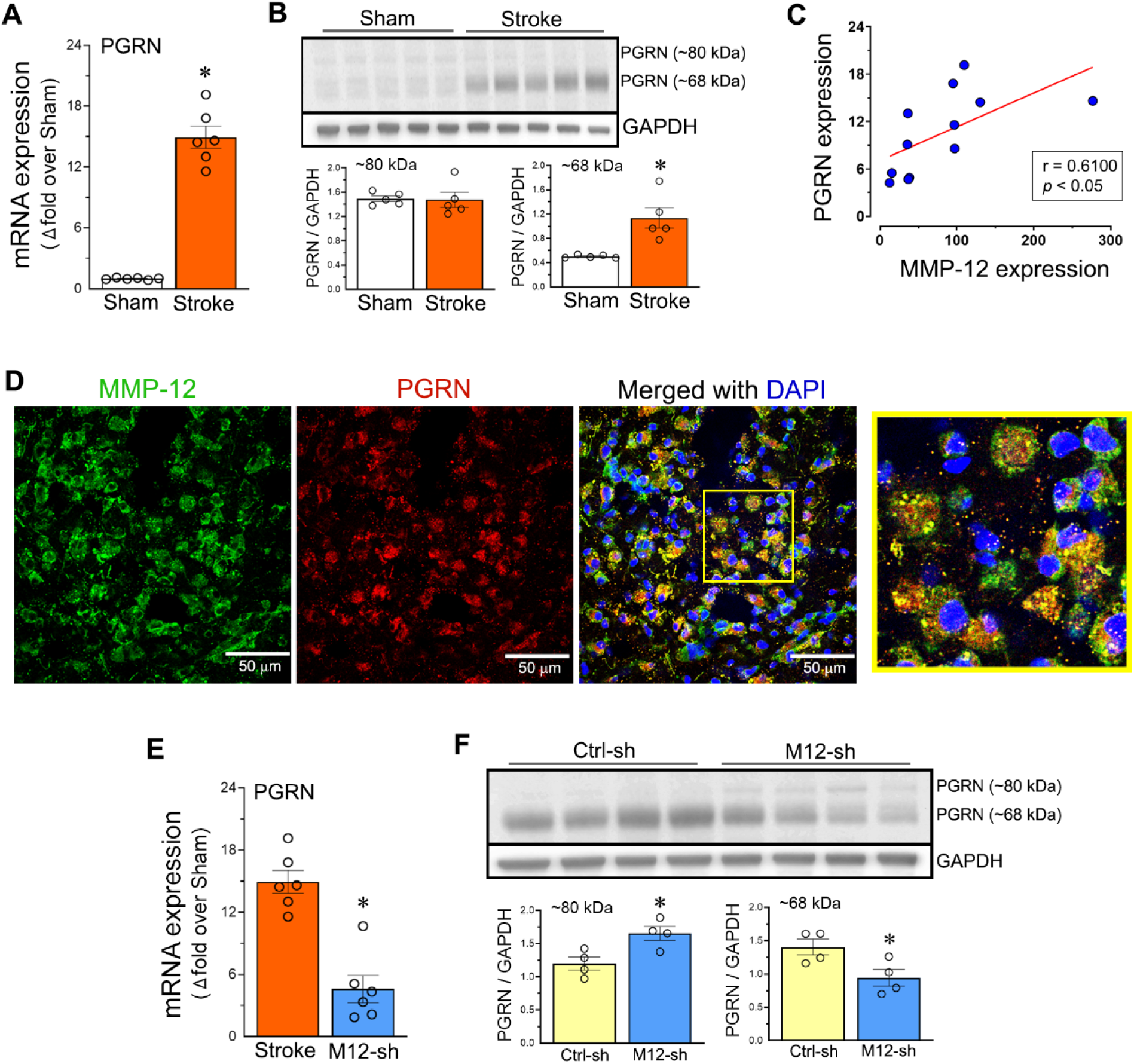
The potential interplay between MMP-12 and PGRN in the ischemic brain. (**A**) Column scatter plot shows the relative PGRN mRNA expression in the ipsilateral hemisphere of rats on post-MCAO day 3. Data are presented as mean ± SEM; n = 6 per group. \**p* < 0.05 vs. Sham group. (**B**) Immunoblots show PGRN protein expression at approximately 80 and 68 kDa, along with GAPDH, in the ipsilateral hemisphere of rats from the Sham and Stroke groups on post-MCAO day 3. Column scatter plot shows the expression of the ∼80- and ∼68-kDa PGRN bands normalized to GAPDH. n = 5 per group. \**p* < 0.05 vs. Sham group. (**C**) Scatter plot shows a significant positive Pearson correlation between relative MMP-12 mRNA expression and relative PGRN mRNA expression in the ipsilateral hemisphere of rats on post-MCAO day 3. n = 12. (**D**) Representative double-labeled immunofluorescence images showing MMP-12 protein expression (green), PGRN protein expression (red), and their colocalization in the peri-infarct cortex of the ipsilateral hemisphere of rats from the Stroke group on post-MCAO day 3. Nuclei (blue) were stained with DAPI. n = 5. (**E**) Column scatter plot shows the relative PGRN mRNA expression in the ipsilateral hemisphere of rats from the Stroke and M12-sh (stroke + MMP-12 shRNA treatment) groups on post-MCAO day 3. n = 6 per group. \**p* < 0.05 vs. Stroke group. (**F**) Immunoblots show PGRN protein expression at approximately 80 and 68 kDa, along with GAPDH, in the ipsilateral hemisphere of rats from the Ctrl-sh (stroke + control shRNA treatment) and M12-sh groups on post-MCAO day 3. Column scatter plot shows the expression of the ∼80- and ∼68-kDa PGRN bands normalized to GAPDH. n = 4 per group. \**p* < 0.05 vs. Ctrl-sh group.

PGRN mRNA expression was positively correlated with MMP-12 mRNA expression (Fig. 7C). Furthermore, PGRN and MMP-12 proteins colocalized in the ischemic brain (Fig. 7D). MMP-12 shRNA treatment significantly attenuated PGRN mRNA expression compared with the Stroke group (Fig. 7E). While the ∼68-kDa PGRN band significantly decreased, the ∼80-kDa band significantly increased in rats in the M12-sh group compared with those in the Ctrl-sh group (Fig. 7F).

## Discussion

This study examined the role of increased MMP-12 expression in acute neuroinflammation following cerebral I/R as well as its relationship with other MMPs, tPA, and PGRN. We observed that MMP-12 exhibited the highest induction in the ischemic brain among the 22 MMPs examined, corroborating our previous findings. In addition, we observed increased expression of several other MMPs (MMP-19, MMP-20, MMP-23, MMP-24, and MMP-27) that had not previously been reported to be upregulated following cerebral ischemia.^2^ MMP-12 is a 54-kDa proenzyme (zymogen containing propeptide, catalytic, and hemopexin domains) that is processed into a 45-kDa form (an intermediate active form containing the catalytic and hemopexin domains) and subsequently into a 22-kDa form (the fully active catalytic domain) and a 23-kDa form (the hemopexin domain).^20^ Although three MMP-12 bands (∼54, ∼45, and ∼23 kDa) were detected in the ischemic brain, the 23-kDa band showed the most pronounced differences in expression among the experimental groups and the greatest consistency within groups. Similar to our previous findings, we observed that MMP-12 shRNA treatment reduced MMP-12 protein expression following cerebral I/R.^2,10^

Although inflammatory responses may initially help contain ischemic tissue injury and aid in tissue repair, excessive neuroinflammation during the acute phase following cerebral I/R exacerbates brain damage.^21^ Neuroinflammation is characterized by BBB disruption, astrogliosis, microgliosis, infiltration of peripheral immune cells such as monocytes and neutrophils, and increased expression of astrocyte A1- and microglia/macrophage M1-type markers, including chemokines and cytokines.^22–24^ We observed a substantial upregulation of IFNγ, a proinflammatory cytokine associated with microglia/macrophage activation and M1 polarization. Consistent with this finding, M1-type marker expression was significantly increased in the ischemic brain. MMP-12 expression was positively correlated with the expression of the astrocyte-associated markers GFAP and vimentin, the proinflammatory A1-type markers C3 and GBP2, and the M1-type markers CCL2, CXCL1, and IL-6. Furthermore, MMP-12 suppression reduced the mRNA expression of GFAP, vimentin, C3, GBP2, CD68, TLR2, TLR4, IL-1β, and IL-6, as well as the protein expression of the proinflammatory cytokines IL-1β, IL-6, and TNFα. The decreased expression of M1 markers following MMP-12 suppression may be attributable, at least in part, to reduced IFNγ protein expression. Collectively, these findings support a role for MMP-12 in exacerbating neuroinflammation following transient focal cerebral ischemia.

Another hallmark of neuroinflammation in the ischemic brain is gliosis. Reactive gliosis observed in animal stroke models reflects cellular responses that occur in the human brain following ischemic stroke.^25^ The altered astrocyte morphology and reduced GFAP expression following MMP-12 suppression suggest that MMP-12 contributes to neuroinflammation by promoting reactive astrogliosis. Similarly, MMP-12 suppression resulted in an intermediate microglia/macrophage morphology with thinner processes and smaller cell bodies than those observed in untreated rats following transient cerebral ischemia, consistent with attenuated cellular activation and microgliosis. These MMP-12 suppression-associated changes in astrocyte and microglia/macrophage activation are consistent with the reduced expression of proinflammatory A1- and M1-type markers, suggesting an overall attenuation of the inflammatory response.

In contrast to the proinflammatory M1-type response, the increased expression of M2-type markers following transient focal cerebral ischemia might represent an endogenous protective response aimed at limiting ischemic brain injury and promoting tissue repair. Interestingly, MMP-12 mRNA expression was positively correlated with M2-type marker expression. This association could reflect MMP-12 expression by M2-type microglia/macrophages or, alternatively, a relationship between elevated MMP-12 levels and the induction of M2-type marker expression in the ischemic brain. The former possibility is supported by previous evidence demonstrating MMP-12 expression in both M1- and M2-type macrophages.^26^

It was recently reported that MMP-12 suppression upregulates M2-type marker expression and may shift microglia/macrophage polarization toward an M2 phenotype.^27^ Therefore, we anticipated that reducing MMP-12 expression would enhance M2-type marker expression following transient focal cerebral ischemia. In contrast, MMP-12 suppression significantly decreased the mRNA expression of the M2-type markers CD163, CD206, Arg1, IL-10, and TGFβ. This finding, together with the MMP-12 shRNA-mediated reduction in several M1-type proinflammatory markers, suggests that MMP-12 suppression attenuates the overall activation of microglia/macrophages, rather than simply promoting a phenotypic shift from M1 to M2. This interpretation is also consistent with the non-significant trend toward reduced Iba1 protein expression following MMP-12 shRNA treatment. Thus, the inhibitory effect of MMP-12 suppression on microglia/macrophage activation may predominate over any potential shift in polarization toward an M2 phenotype following focal cerebral ischemia.

BBB disruption following transient focal cerebral ischemia leads to extensive infiltration of neutrophils and monocytes exacerbating neuroinflammation.^28^ The chemokine CCL2, also referred to as MCP-1, binds to CCR2, a receptor expressed on monocytes, and facilitates their extravasation into the ischemic brain.^29^ Similarly, the chemoattractant protein CXCL1 binds to CXCR2, a receptor expressed on neutrophils, and promotes neutrophil recruitment into the ischemic brain.^30^ Consistent with these processes, we observed substantial increases in the expression of CCL2 and CXCL1 in the ischemic brain, suggesting increased recruitment of monocytes and neutrophils. Significant upregulation of the monocyte/macrophage- and neutrophil-associated markers CCR2, CXCR2, ELANE, and MPO further supports the involvement of these cells in the post-ischemic inflammatory response. MMP-12 mRNA expression was positively correlated with monocyte/macrophage- and neutrophil-associated markers, suggesting that infiltrating monocytes and neutrophils may represent additional cellular sources of MMP-12 in the ischemic brain. These results are consistent with previous reports that macrophages and neutrophils express MMP-12, which was initially referred to as macrophage metalloelastase.^31,32^ MPO, a well-known proinflammatory enzyme expressed predominantly by neutrophils, was significantly reduced following MMP-12 suppression. Given our previous finding that MMP-12 suppression attenuates BBB disruption following transient focal cerebral ischemia, preservation of BBB integrity may limit the infiltration of leukocytes into the ischemic brain.^10^ Reduced leukocyte infiltration may, in turn, decrease the contribution of leukocyte-derived MMPs and other proinflammatory mediators, including MPO, to secondary brain injury. Previous studies have demonstrated that infiltrating monocytes and neutrophils can express a variety of MMPs.^33,34^ Consistent with this possibility, MMP-12 suppression significantly reduced the mRNA expression of numerous other MMPs in the ischemic brain. Because multiple MMPs can contribute to BBB disruption following transient focal cerebral ischemia, leukocyte-derived MMPs may further compromise BBB integrity and facilitate additional infiltration, thereby amplifying neuroinflammation and secondary brain injury.^35^ Thus, by attenuating BBB disruption, MMP-12 suppression might interrupt this cycle, limiting leukocyte infiltration and the subsequent accumulation of additional MMPs and proinflammatory mediators in the ischemic brain.

Numerous studies have reported an increase in both the expression and activity of endogenous tPA in the brain following transient focal cerebral ischemia.^10,12,36–38^ Elevated endogenous tPA in the ischemic brain may increase the expression of several MMPs, including MMP-12, through BBB disruption and may also promote the activation of multiple MMPs, including MMP-12, through plasmin formation.^12,39–46^ Computational modeling of protein-protein interactions suggested a potential physical interaction between MMP-12 and tPA.^12^ This prediction is further supported by the prominent colocalization of MMP-12 and tPA observed in the ischemic brain in the present study. Furthermore, the co-immunoprecipitation of tPA with MMP-12 using an anti-MMP-12 antibody provides additional evidence of an association between these proteins. Increased endogenous tPA levels might indirectly upregulate MMP-12 expression through BBB disruption and subsequent monocyte and neutrophil infiltration, potentially introducing additional cellular sources of MMP-12 into the ischemic brain. However, the functional consequences of the potential physical interaction between MMP-12 and tPA remain unknown. MMP-12 suppression significantly reduced endogenous tPA protein expression on post-MCAO day 3, consistent with our previous findings on post-MCAO day 1.^10^ Previous studies have demonstrated that tPA contributes to chemokine activation, recruitment of infiltrating monocytes, microglial activation, and increased expression of proinflammatory microglia/macrophage M1-type markers.^12,47–51^ Thus, the attenuation of neuroinflammation following MMP-12 suppression may be mediated, at least in part, by reduced endogenous tPA expression in the ischemic brain.

PGRN is known for its anti-inflammatory and neuroprotective properties.^52–56^ The significant upregulation of PGRN mRNA and the ∼68-kDa PGRN protein observed in the present study is consistent with a recent report demonstrating that PGRN protein expression in the ischemic brain of mice peaked on post-MCAO day 3.^57^ A variety of cell types in the ischemic brain can express PGRN; however, microglia/macrophages are considered a major source.^57^ In the ischemic brain, PGRN protects the BBB, attenuates brain hemorrhage, neuroinflammation, and ferroptosis, reduces infarct volume, and promotes functional recovery and neurogenesis.^57–60^ Thus, increased PGRN expression in the ischemic brain might represent an endogenous protective response to the injury following transient focal cerebral ischemia. The prominent colocalization of MMP-12 and PGRN observed in the current study demonstrates their spatial association in the peri-infarct cortex and raises the possibility of an interaction between them. MMP-12 suppression significantly increased the ∼80-kDa PGRN band, potentially representing a mature, glycosylated form while decreasing the ∼68-kDa band, potentially representing a less extensively glycosylated or unmodified form, suggesting that reducing MMP-12 expression may alter PGRN processing in the ischemic brain. Reduced proteolytic cleavage of anti-inflammatory PGRN following MMP-12 suppression could contribute to the attenuation of neuroinflammation observed in the present study. This possibility is consistent with a previous report implicating MMP-12 in the proteolytic cleavage of PGRN into proinflammatory granulins that promote cytokine expression.^16^ Interestingly, MMP-12 suppression also significantly reduced PGRN mRNA expression. Given the proposed compensatory upregulation of PGRN following transient focal cerebral ischemia, this reduction might reflect a diminished need for PGRN production as neuroinflammation is attenuated following MMP-12 suppression. Together, these findings support a proposed model linking MMP-12-mediated PGRN processing with neuroinflammation following cerebral ischemia and reperfusion (Supplementary Fig. 10).

The exclusive use of young male rats to evaluate the role of MMP-12 in post-ischemic neuroinflammation is a limitation of this study. However, this approach was intended to reduce biological variability during the initial assessment of treatment efficacy and elucidation of the underlying molecular mechanisms. This is consistent with the recommendations of the Stroke Therapy Academic Industry Roundtable (STAIR), which recommended initial evaluation in young, healthy male animals before extending investigations to females, aged animals, and those with comorbid conditions.^61^ Although the effects of MMP-12 suppression on neuroinflammation were not evaluated in female rats in the present study, our previous finding that MMP-12 suppression enhanced functional recovery following transient focal cerebral ischemia in both male and female rats suggests that this benefit is not restricted to one sex.^11^

In summary, MMP-12 exhibited the greatest upregulation among the MMPs examined in the ischemic brain, and its expression was positively correlated with neurological deficit severity following transient focal cerebral ischemia. MMP-12 expression was positively correlated with astrocyte-associated and proinflammatory A1-type markers, microglia/macrophage M1-type markers, as well as monocyte/macrophage- and neutrophil-associated markers in the ischemic brain. MMP-12 suppression altered the morphology of astrocytes and microglia/macrophages, consistent with reduced cellular activation and gliosis, and reduced the expression of multiple astrocyte and proinflammatory microglia/macrophage markers, as well as the neutrophil marker MPO. Furthermore, MMP-12 suppression reduced the expression of several other MMPs, as well as endogenous tPA, and altered PGRN expression in a manner suggesting differential processing. Collectively, these findings suggest that MMP-12 contributes to acute post-ischemic neuroinflammation, and that its suppression attenuates neuroinflammation, possibly through changes in other MMPs, tPA, and PGRN.

## Author statement

KKV, JDK, and RV contributed to the conception, funding acquisition, and oversight of the project. KKV and SRC performed stroke surgeries. CAF, SRC, and IMB were involved in post-operative animal care. SRC, CAF, IMB, PU, SNJ, VV, JN, VS, ATV, KRP, KAR, and JB were involved in methodology and acquisition of data. KKV and SRC performed data analysis and interpretation. KKV drafted the manuscript. CAF and RV edited the manuscript. All the authors reviewed and approved the final manuscript.

## Funding

This work was supported by the National Institutes of Health (R01NS102573 to KKV and R35NS132184 to RV). KKV is the recipient of the Transformational Project Award (23TPA1061995) from the American Heart Association, while RV is the recipient of a Research Career Scientist Award (IK6BX005690) from the U.S. Department of Veterans Affairs.

## Notes

### Competing Interest Statement

The authors have declared no competing interest.

