## Supplemental Material for "MMP-12 exacerbates neuroinflammation in the ischemic brain: possible interactions with tPA and progranulin"

### Supplementary Material

#### Methods

##### *Real-time PCR*

Brain tissue from rats in the appropriate experimental groups was collected on post-MCAO day 3. Under deep isoflurane anesthesia, rats were perfused intracardially with ice-cold 1× PBS, and the brains were rapidly removed. Total RNA from the entire ipsilateral cerebral hemisphere was extracted using TRIzol reagent (Thermo Fisher Scientific, Waltham, MA, USA). One microgram of total RNA from each sample was reverse transcribed into cDNA using the iScript cDNA Synthesis Kit (Bio-Rad Laboratories, Hercules, CA, USA). For each diluted (1:10) cDNA sample, reactions were assembled using iTaq Universal SYBR Green Supermix (Bio-Rad Laboratories, Hercules, CA, USA). Forward and reverse primers for the target genes (Integrated DNA Technologies, Coralville, IA, USA) were diluted 1:10 in nuclease-free water. Primer sequences for the target genes are listed in Supplementary Table 2. The specificity of the primers for each target gene was evaluated by melting-curve analysis and agarose gel electrophoresis of PCR products. Melting curves showed a single peak, and agarose gel electrophoresis yielded a single PCR product for each gene, supporting the specificity of the primers selected for qPCR analysis.

Real-time PCR reactions were performed in triplicate using the following thermal cycling conditions: initial denaturation at 95 ˚C for 5 min; 40 cycles of 95 ˚C for 30 s, 58-62 ˚C for 30 s, and 72 ˚C for 30 s; and a final extension at 72 ˚C for 5 min. Reactions were run on an iCycler iQ Multi-Color Real-Time PCR Detection System (Bio-Rad Laboratories, Hercules, CA, USA). Data were collected using iCycler iQ software (Bio-Rad Laboratories, Hercules, CA, USA) and expressed as cycle threshold (Ct) values. For each sample, the mean Ct value from triplicate reactions after exclusion of any outlier was used for analysis. *18S* rRNA served as the internal reference. Relative target gene expression was normalized to *18S* rRNA, and fold change (test vs. control) was calculated as 2^(ΔCt control)/2^(ΔCt test), where ΔCt = Ct (target) - Ct (*18S* rRNA).

***Immunoprecipitation***

Immunoprecipitation of MMP-12 from the ipsilateral cerebral hemisphere of rats from the Stroke group was performed using the Dynabeads™ Protein A Immunoprecipitation Kit (Thermo Fisher Scientific, Waltham, MA, USA). Briefly, 50 µL (1.5 mg) of Dynabeads™ Protein A magnetic beads were resuspended, transferred to a tube, and isolated using a magnetic bead separation rack (Millipore Sigma, Burlington, MA, USA). The magnetic beads were incubated under rotation with 10 µg of anti-MMP-12 antibody (Supplementary Table 3) diluted in 200 µL of antibody (Ab) Binding and Washing Buffer for 20 min. The magnetic bead–antibody complex was separated using the magnetic bead separation rack, the supernatant was removed, and the beads were washed once by gentle pipetting with 200 µL of Ab Binding and Washing Buffer. For capture of the MMP-12 antigen from ipsilateral cerebral hemisphere tissue lysates from rats in the Stroke group, 1 mg of protein lysate from each rat, diluted in 500 µL of lysis buffer containing protease and phosphatase inhibitors, was added to the magnetic bead–antibody complex. The mixture was incubated under continuous rotation overnight at 4 °C. Following incubation, the supernatant (flow-through) was collected for downstream analysis. The magnetic bead–antibody–antigen complex was washed three times with washing buffer, and the magnetic beads were collected between each wash step using the magnetic bead separation rack. To minimize coelution of proteins bound to the tube wall, the complex was resuspended in

100 μL of washing buffer, and the magnetic bead suspension was transferred to a clean microcentrifuge tube prior to final magnetic bead separation. The bound target proteins were eluted under denaturing and reducing conditions by resuspending the magnetic beads in 20 μL of elution buffer, 5 μL of NuPAGE LDS sample buffer, and 0.5 μL of β-mercaptoethanol. The tube containing the sample mixture was heated at 70 °C for 10 min in a heating block. The tube was then placed on the magnetic bead separation rack, and the supernatant containing the eluted proteins was collected. Both the collected flow-through fraction and the final eluate from each sample were subjected to immunoblot analysis, as described in the previous section, using a rabbit monoclonal anti-tPA primary antibody and an HRP-conjugated goat anti-rabbit secondary antibody (Supplementary Table 3).

***Tissue sectioning and immunofluorescence***

On post-MCAO day 3, rats were deeply anesthetized and perfused transcardially with 1× PBS, followed by 10% neutral buffered formalin (Fisher Scientific, Hampton, NH, USA). Brains were removed, postfixed in the same formalin solution for 24 h, and cryoprotected in 30% sucrose in 1× PBS until they sank. The brains were then embedded in Tissue-Tek® O.C.T. Compound (Electron Microscopy Sciences, Hatfield, PA, USA) and sectioned using a Leica CM1950 cryostat (Leica Microsystems, Deerfield, IL, USA). Frozen coronal sections (40 μm thick) were collected through the striatum and overlying cortex at the level of the anterior commissure. Following sectioning, free-floating sections were collected in 24-well polystyrene plates containing 0.02% sodium azide and stored at 4 °C until use.

For each animal, a single coronal brain section was processed for immunofluorescence. Sections were mounted onto microscope slides from 50 mM phosphate buffer (pH 7.4), dried overnight at 37 °C, and maintained at room temperature for an additional day prior to processing. Slide-mounted sections were subjected to heat-induced epitope retrieval in 10 mM citric acid (pH 6.0) at 90 °C for 30 min in a water bath. Following rinses with 1× PBS, sections were permeabilized with 0.3% Triton X-100 in 1× PBS and blocked with 5% normal goat or donkey serum containing 0.3% Triton X-100 in 1× PBS. Following blocking, sections were incubated overnight at 4 °C with the appropriate primary antibodies (Supplementary Table 3) at the specified dilutions in 3% normal goat or donkey serum containing 0.3% Triton X-100 in 1× PBS. The following day, sections were rinsed with 1× PBS and incubated with the appropriate Alexa Fluor 488- or Alexa Fluor 594-conjugated IgG secondary antibodies (Supplementary Table 3) at the specified dilutions in 1× PBS. Secondary antibody incubation was performed for 3 h in the dark, consisting of 1 h at room temperature followed by 2 h at 4 °C. Sections were subsequently counterstained with 4′,6-diamidino-2-phenylindole (DAPI; 1 μg/mL in 1× PBS) for 15 min at room temperature in the dark to visualize cell nuclei. Following rinses, sections were incubated for 10 min in an aqueous solution containing 10 mM copper sulfate and 50 mM ammonium chloride to quench lipofuscin autofluorescence. After additional rinses, slides were coverslipped using Immu-Mount™ (Epredia, Netherlands) and allowed to dry overnight at room temperature, and subsequently stored at 4 °C, with slides protected from light throughout.

Images were acquired using a 40× objective on an Olympus FluoView FV3000 confocal laser scanning microscope. For each animal, one field of view within the peri-infarct region was imaged at a consistent anatomical location across animals. Identical imaging and channel settings were used for all samples to permit direct comparison of fluorescence staining among experimental groups. Quantification of immunofluorescence was performed using NIH ImageJ software (version 1.53e). Images were converted to 8-bit grayscale, and, for each antigen or marker, an optimal threshold was empirically established and subsequently held constant for analysis of all corresponding experimental groups. Integrated density (arbitrary units) was used as the quantitative measure of immunofluorescence. Astrocyte and microglia/macrophage morphology and co-labeling of selected markers were also visually examined.

***Statistical analysis***

Statistical analyses were performed using GraphPad Prism 10.4.2 for Windows (GraphPad Software, San Diego, CA, USA). For real-time PCR data, statistical analyses were performed on ΔCt values. Quantitative data were tested for normality (Shapiro-Wilk and Kolmogorov-Smirnov tests) and homogeneity of variance (F-test, Bartlett's test, or Brown-Forsythe test). Based on the outcomes of these tests, appropriate statistical tests were applied.

For comparisons involving only two groups, the Mann-Whitney test was used when normality was not met; for normally distributed data, a two-tailed unpaired t-test was used, with Welch's correction when variances were unequal. Because multiple hypothesis testing increases the risk of false-positive findings, the false discovery rate (FDR) correction was applied and controlled at $Q = 0.05$ using the two-stage linear step-up procedure of Benjamini, Krieger, and Yekutieli, applied separately for each family of genes (MMPs, A1 markers, M1 markers, M2 markers, and leukocyte markers). For multiple-group comparisons, one-way ANOVA followed by Tukey's multiple comparisons test was used when variances were equal and Brown-Forsythe or Welch's ANOVA followed by Dunnett's T3 multiple comparisons test was used when variances were unequal. For comparisons involving two different variables, two-way ANOVA followed by Sidak's multiple comparisons test was used. For assessing the relationship between two variables, the Pearson correlation test was used when normality was met, whereas the Spearman correlation test was used when normality was not met.

Differences between groups were considered significant at $p < 0.05$ or FDR-adjusted $p < 0.05$, as appropriate. Supplementary Table 4 summarizes the statistical analyses, including the tests applied, test statistics (t ratios and t- and F-values), and exact *p*-values.

### Supplementary Figure 1

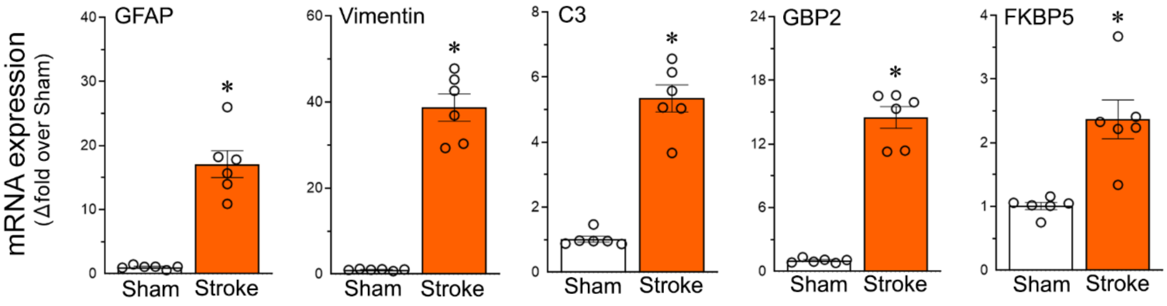


**Supplementary Fig. 1. Astrocyte marker expression increases in the brain following cerebral I/R.** Column scatter plots show the relative mRNA expression of astrocyte-associated markers GFAP and vimentin and proinflammatory A1-type markers C3, GBP2, and FKBP5 in the ipsilateral hemisphere of rats on post-MCAO day 3. Data are presented as mean ± SEM; n = 6 per group. *$p < 0.05$ vs. Sham group.

**Supplementary Figure 2**

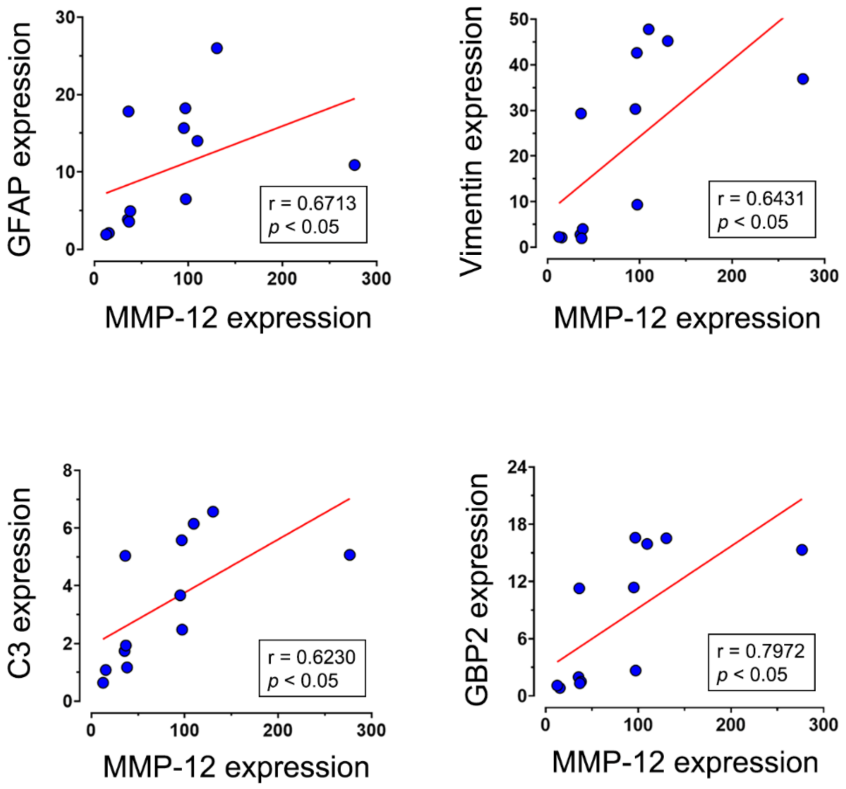


**Supplementary Fig. 2. Association between MMP-12 and astrocyte markers in the brain following cerebral I/R.** Scatter plots show significant positive correlations (Pearson or Spearman correlation, as appropriate) between the relative mRNA expression (fold change over Sham) of MMP-12 and astrocyte-associated markers GFAP and vimentin and proinflammatory A1-type markers C3 and GBP2 in the ipsilateral hemisphere of rats on post-MCAO day 3. n = 12.

**Supplementary Figure 3**

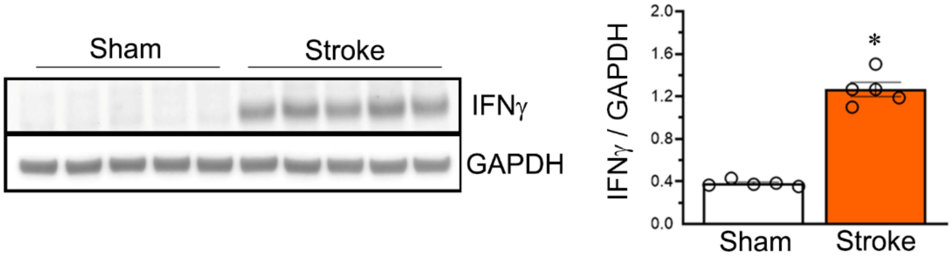


**Supplementary Fig. 3. IFNγ expression increases in the brain following cerebral I/R.** Immunoblot shows the protein expression of IFNγ and GAPDH in the ipsilateral hemisphere of rats on post-MCAO day 3. Column scatter plot shows IFNγ protein expression normalized to GAPDH. Data are presented as mean ± SEM; n = 5 per group. *$p < 0.05$ vs. Sham group.

#### Supplementary Figure 4

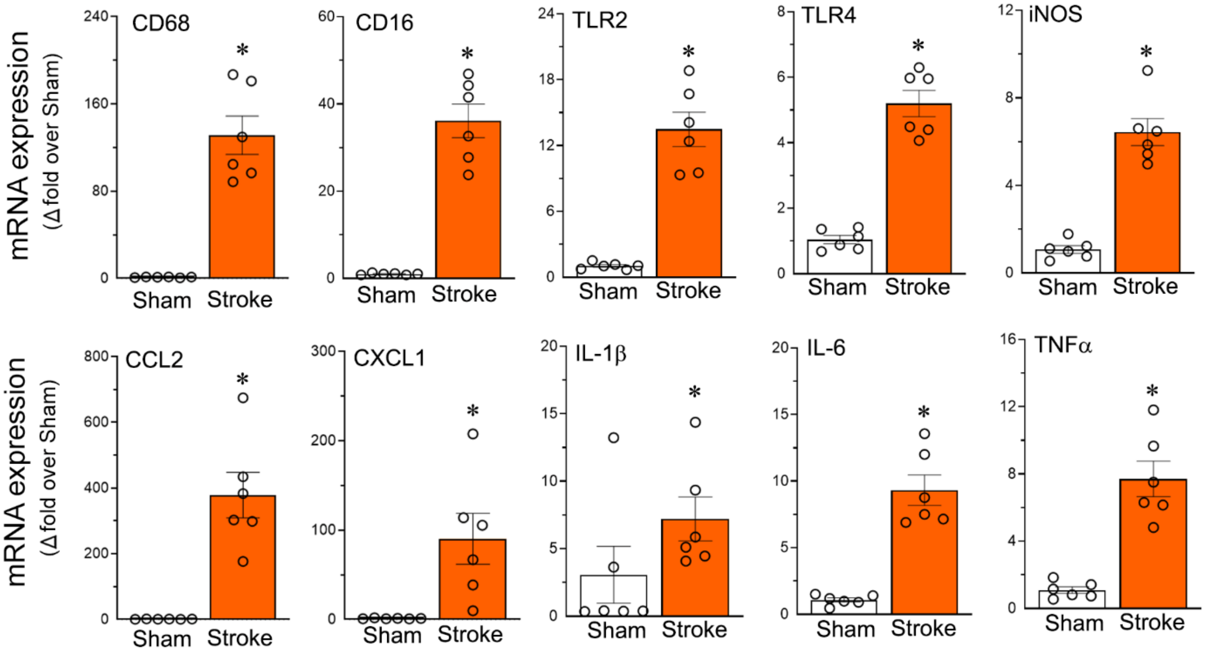


**Supplementary Fig. 4. Microglia/macrophage M1-type proinflammatory marker expression increases in the brain following cerebral I/R.** Column scatter plots show the relative mRNA expression of M1-type proinflammatory markers CD68, CD16, TLR2, TLR4, iNOS, CCL2, CXCL1, IL-1β, IL-6, and TNFα in the ipsilateral hemisphere of rats on post-MCAO day 3. Data are presented as mean ± SEM; n = 6 per group. *$p < 0.05$ vs. Sham group.

**Supplementary Figure 5**

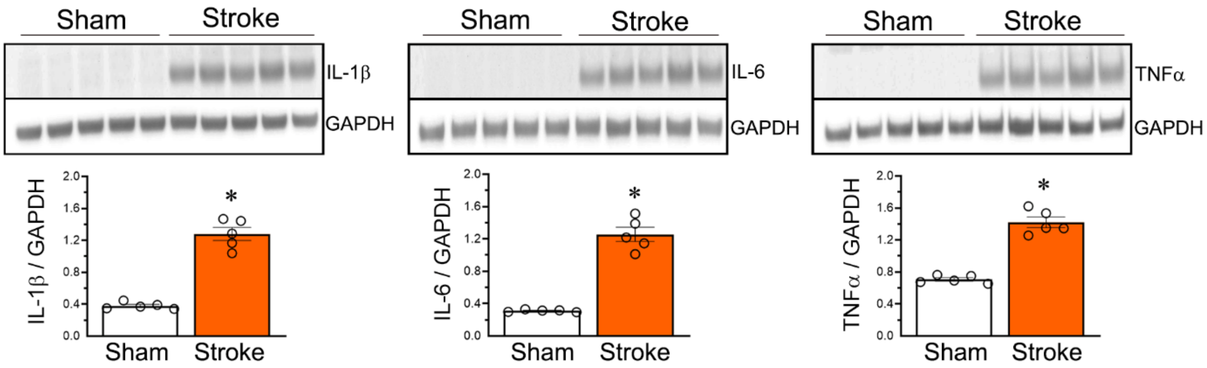


**Supplementary Fig. 5. The protein expression of proinflammatory cytokines increases in the brain following cerebral I/R.** Immunoblots show the protein expression of the proinflammatory cytokines IL-1β, IL-6, and TNFα and GAPDH in the ipsilateral hemisphere of rats from the Sham and Stroke groups on post-MCAO day 3. Column scatter plots show IL-1β, IL-6, and TNFα protein expression normalized to GAPDH. Data are presented as mean ± SEM; n = 5 per group. *$p < 0.05$ vs. Sham group.

**Supplementary Figure 6**

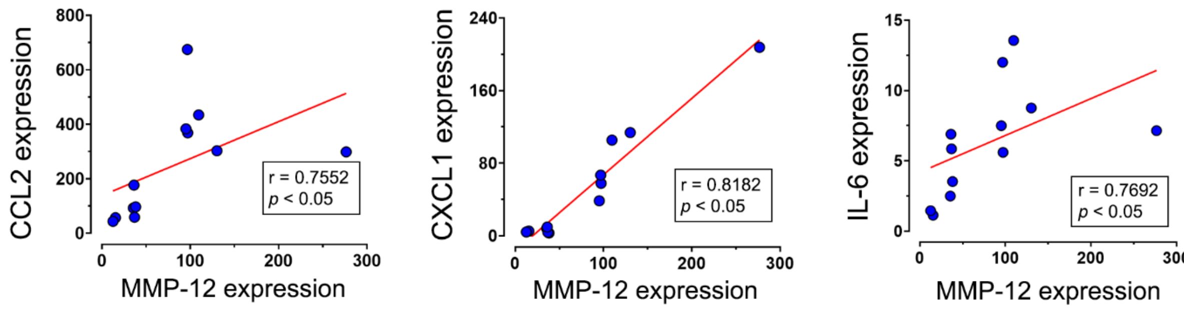


**Supplementary Fig. 6. MMP-12 expression positively correlates with the expression of microglia/macrophage M1-type proinflammatory markers in the brain following cerebral I/R.** Scatter plots show significant positive correlations (Pearson or Spearman correlation, as appropriate) between relative mRNA expression (fold change over Sham) of MMP-12 and the M1-type proinflammatory markers CCL2, CXCL1, and IL-6 in the ipsilateral hemisphere of rats on post-MCAO day 3. n = 12.

#### Supplementary Figure 7

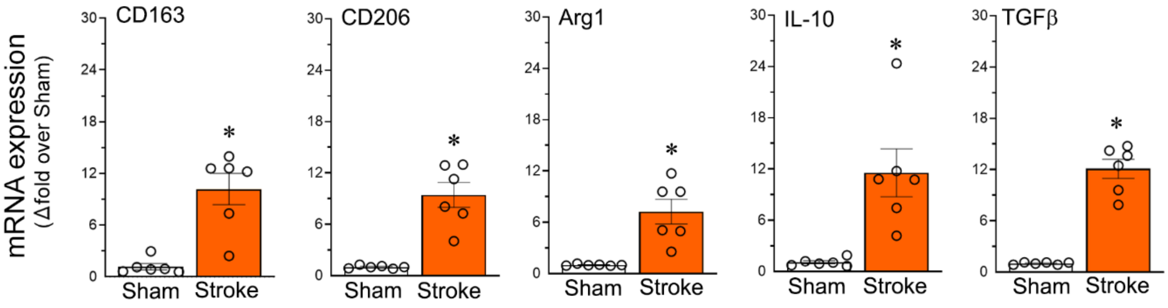


**Supplementary Fig. 7. Microglia/macrophage M2-type anti-inflammatory marker expression increases in the brain following cerebral I/R.** Column scatter plots show the relative mRNA expression of M2-type anti-inflammatory markers CD163, CD206, Arg1, IL-10, and TGFβ in the ipsilateral hemisphere of rats on post-MCAO day 3. Data are presented as mean ± SEM; n = 6 per group. **p* < 0.05 vs. Sham group.

#### Supplementary Figure 8

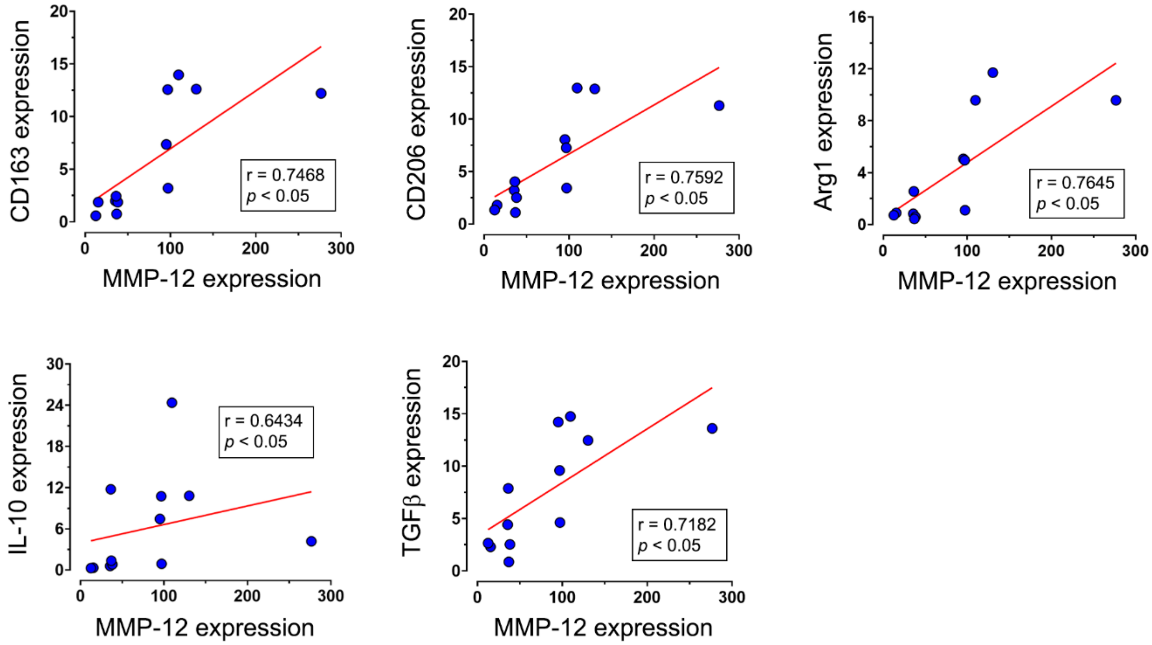


**Supplementary Fig. 8. MMP-12 expression positively correlates with the expression of microglia/macrophage M2-type anti-inflammatory markers in the brain following cerebral I/R.** Scatter plots show significant positive correlations (Pearson or Spearman correlation, as appropriate) between relative mRNA expression (fold change over Sham) of MMP-12 and M2-type anti-inflammatory markers CD163, CD206, Arg1, IL-10, and TGFβ in the ipsilateral hemisphere of rats on post-MCAO day 3. n = 12.

**Supplementary Figure 9**

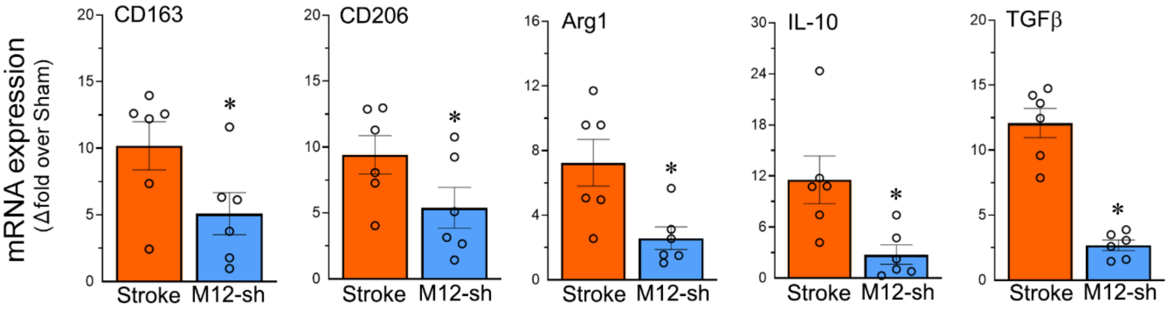


**Supplementary Fig. 9. MMP-12 inhibition following cerebral I/R reduces the expression of microglia/macrophage M2-type markers in the ischemic brain.** Column scatter plots show the relative mRNA expression of M2-type anti-inflammatory markers CD163, CD206, Arg1, IL-10, and TGFβ in the ipsilateral hemisphere of rats from the Stroke and M12-sh (stroke + MMP-12 shRNA treatment) groups on post-MCAO day 3. Data are presented as mean ± SEM; n = 6 per group. **p* < 0.05 vs. Stroke group.

**Supplementary Figure 10**

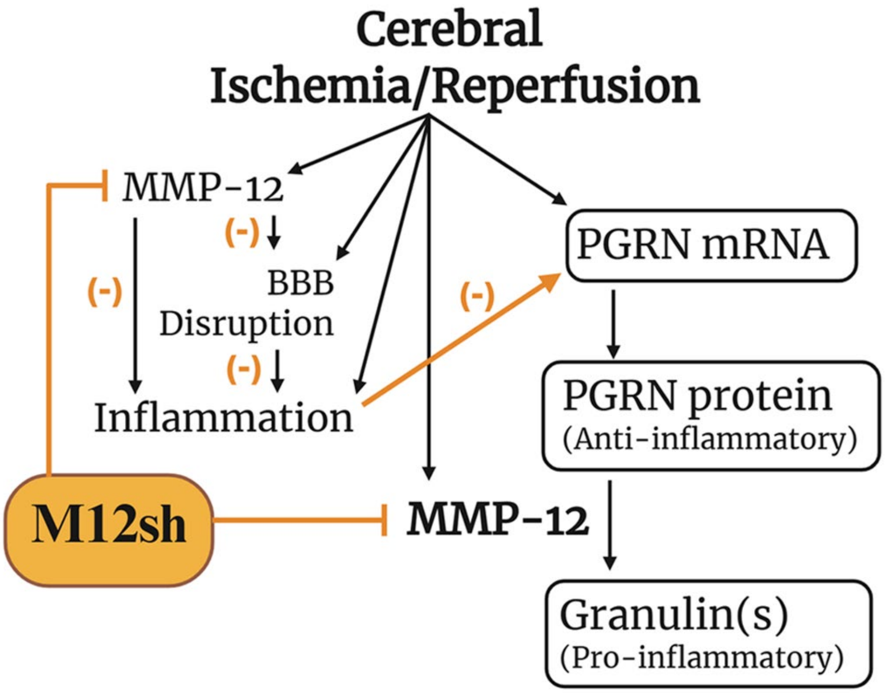


**Supplementary Fig. 10. Proposed model of MMP-12-mediated PGRN processing and inflammation following cerebral ischemia/reperfusion (I/R) and the potential effects of MMP-12 shRNA treatment.** Cerebral I/R induces upregulation of MMP-12 and progranulin (PGRN) in the ischemic brain. Increased MMP-12 contributes to blood-brain barrier (BBB) disruption and inflammation and may proteolytically cleave anti-inflammatory PGRN protein into proinflammatory granulins.[10,16] MMP-12 suppression by MMP-12 shRNA treatment may limit the proteolytic cleavage of PGRN into granulins, reduce BBB disruption,[10] attenuate downstream inflammatory signaling, and thereby indirectly decrease PGRN mRNA production. This figure was created with biorender.com under BioRender's Academic Institutional Subscription.

**Supplementary Table 1.** Experimental groups, description, and animal numbers.

| Group Name | Group Description | Number of animals | | | | |
|---|---|---|---|---|---|---|
| | | PCR | Immunoblot | IF | Excluded | Total |
| Sham | Rats subjected to MCAO surgery (no suture inserted). | 6 | 5 | 6 | 0 | 17 |
| Stroke | Rats subjected to 2-h MCAO but received no treatment | 12 | 5 | 6 | 9 | 32 |
| Ctrl-sh (Stroke + Control shRNA treatment) | Rats subjected to 2-h MCAO and received control shRNA treatment immediately (within 30 min) after reperfusion. | – | 4 | – | 3 | 7 |
| M12-sh (Stroke + MMP-12 shRNA treatment) | Rats subjected to 2-h MCAO and received MMP-12 shRNA treatment immediately (within 30 min) after reperfusion. | 6 | 4 | 6 | 7 | 23 |
| | | | | | TOTAL | 79 |

MCAO, middle cerebral artery occlusion; IF, immunofluorescence; PCR, polymerase chain reaction

**Supplementary Table 2.** Rat primers used for real time PCR analysis.

| Gene | NCBI Reference Sequence | Primer Sequence | |
|---|---|---|---|
| | | Forward (5’ - 3’) | Reverse (5’ - 3’) |
| MMP-1 | NM_001134530 | cagcagttatttgggctgaaag | tttggtccaacgaggattgt |
| MMP-2 | NM_031054 | gctgatactgacactggtactg | tgtcactgtccgccaaataa |
| MMP-3 | NM_133523 | ggaccagggattaatggagatg | agcattggctgagtgaaaga |
| MMP-7 | NM_012864 | gagtgccagatgttgcagaa | ctgcagtcccccaactaa |
| MMP-8 | NM_022221 | tgggctctaagtgcctatga | cgtatctccagcattggttgt |
| MMP-9 | NM_031055 | cactgtaactgggggcaact | cacttcttgtcagcgtcgaa |
| MMP-10 | NM_133514 | tcccaccgtgaagaagattg | cagtctcgggaagcctttat |
| MMP-11 | NM_012980 | gctgagggctatgcctact | gaagcccaggtccacaaata |
| MMP-12 | NM_053963 | gctagaagtaactgggcaact | gagataccgcttcatccatctt |
| MMP-13 | NM_133530 | gccctgatgtttcccatctat | ggtcacacttctctggtgttt |
| MMP-14 | NM_031056 | gtaccccaagtcagctct | cagtgaacgctggcagtaaa |
| MMP-15 | NM_001106168 | aacgtcaggatgggcattt | tgtgtcaatgcggtcatagg |
| MMP-16 | NM_080776 | cagctctggaagaaggtt | gcctgactgcatggtctct |
| MMP-17 | NM_001105925 | ctgtactggcgctatgacga | ctgcctctagctcaccat |
| MMP-19 | NM_001107159 | acttgggcattcccgatatac | acctcttcctcctcatcttct |
| MMP-20 | NM_001106800 | ggtcctcctcgcacaattta | agctgtagtattcctctccca |
| MMP-21 | NM_001106308 | gggtcatgtgagggatcattt | agctgttgcggaagaagtag |
| MMP-23 | NM_053606 | ctgatgcactcacagcaaga | gcccaggatgtacacacaaa |
| MMP-24 | NM_031757 | tgagcaggaggaggagaaata | tcgaaagcctgacgaatagc |
| MMP-25 | NM_001427286 | tggagtatccagtccttctct | tgagaatccacctcttggaac |
| MMP-27 | NM_001106799 | caggcatatctcaaccagttct | tttcccgtcactgtcaatcc |
| MMP-28 | NM_001079888 | gcaaagccaggtcacaaatg | gcgctgacacattactccata |
| GFAP | NM_017009 | gagtggtatcggtccaagttt | ttggcggcgatagtcattag |
| Vimentin | NM_031140 | ccaggacctgctcaatgtaaa | tctctcaggttcagggaagaa |
| C3 | NM_016994 | tcaggtagcgacatggtaga | aacaccatgaggtcgaaagg |
| GBP2 | NM_133624 | tgctaactttgtgggcttct | ttcttcagggtcagcgaatg |
| FKBP5 | NM_001012174 | gggagatggacaccaaagaaa | atctccagccaggacactat |
| CD68 | NM_001031638 | acaaacagtccaggcttctc | acatggctgggaaccattag |
| CD16 | NM_207603 | acttctgcagagggatcattg | cagcaggcagaaagtgatttg |
| TLR2 | NM_198769 | ggaagcaggtgacaaccatt | cgcctaagagcaggatcaac |
| TLR4 | NM_019178 | gtgggtcaaggaccagaaaa | ggctaccacaagcacactga |
| iNOS | NM_012611 | agggccacctttatgtttgt | gctcttgagctggaagaagtaa |
| CCL2 | NM_031530 | gtctcagccagatgcagttaat | gctgctggtgattctcttgt |
| CXCL1 | NM_030845 | gcacccaaaccgaagtcata | gggacaccctttagcatcttt |
| IL-1β | NM_031512 | gctttcgacagtgaggagaat | cgagatgctgctgtgagatt |
| IL-6 | NM_012589 | ccaacttccaatgctctcctaa | ttgccgagtagacctcatagt |
| TNFα | NM_012675 | cagccgatttgccatttcatac | aggtacatgggctcatacca |
| CD163 | NM_001107887 | tgaagtgcctcccaagaatg | ggcaatgagaaggaccaataga |
| CD206 | NM_001106123 | gctgtctgtgcatttccattc | cagcttgtcggtgtcatagt |

| | | | |
|---|---|---|---|
| Arg1 | NM_017134 | ccctgacctatgcgtcattt | cctggtacatctgggaacttt |
| IL-10 | NM_012854 | catggcccagaaatcaagga | agcgtcgcagctgtatcca |
| TGFβ | NM_021578 | ttcagctccacagagaagaac | gtgtccaggctccaaatgta |
| CCR2 | NM_021866 | tgcctctctaccaggaatca | gggcaggatcaaactcaagata |
| CXCR2 | NM_017183 | tcaccatgtggtttctctcac | caccctccacttggatgtatt |
| ELANE | NM_001106767 | ccaaggcattgactccttca | gtcatcatggctgcggataa |
| MPO | NM_001401812 | gctactcctgccaaactgaat | tgcatgttaagggcaggtaag |
| tPA | NM_013151 | ctgtcccagaagccctacag | gccacggtaagtcacacctt |
| PGRN | NM_001145842 | aggttgggaatgtggaatgt | ctctacagcagacaccctttac |
| *18S* rRNA | NR_046237 | acgtctgccctatcaactttc | ttggatgtggtagccgtttc |

PCR, polymerase chain reaction; MMP, matrix metalloproteinase; TLR, toll-like receptor; iNOS, inducible nitric oxide synthase; CCL2, C-C motif chemokine ligand 2; CXCL1, C-X-C motif chemokine ligand 1; IL, interleukin; TNF, tumor necrosis factor; Arg1, arginase 1; TGF, transforming growth factor; CCR2, C-C motif chemokine receptor 2; CXCR2, C-X-C Motif Chemokine Receptor 2; ELANE, elastase, neutrophil expressed; MPO, myeloperoxidase; GFAP, glial fibrillary acidic protein; C3, complement component 3; GBP2, guanylate-binding protein 2; FKBP5, FK506 binding protein 5; tPA, tissue-type plasminogen activator; PGRN, progranulin.

**Supplementary Table 3.** Primary and secondary antibodies for immunoprecipitation, immunoblotting, and immunofluorescence.

| Application | Antibody | Source | Catalog number | Dilution |
|---|---|---|---|---|
| Primary antibodies for immunoprecipitation | MMP-12 – rabbit polyclonal | Bioss | bs-1854R | NA |
| Primary antibodies for immunoblotting | MMP-12 – mouse monoclonal | Santa Cruz Biotechnology | sc-390863 | 1:1,000 |
| | IFN-γ – mouse monoclonal | Santa Cruz Biotechnology | sc-59992 | 1:1,000 |
| | IL-1β – mouse monoclonal | Santa Cruz Biotechnology | sc-52012 | 1:1,000 |
| | IL-6 – mouse monoclonal | Santa Cruz Biotechnology | sc-57315 | 1:1,000 |
| | TNFα – mouse monoclonal | Santa Cruz Biotechnology | sc-52746 | 1:1,000 |
| | MPO – mouse monoclonal | Novus Biologicals | NBP1-51148 | 1:1,000 |
| | tPA – rabbit monoclonal | Novus Biologicals | NBP3-16356 | 1:1,000 |
| | PGRN – sheep polyclonal | R&D Systems | AF2557 | 1:1,000 |
| | GAPDH – mouse monoclonal | Santa Cruz Biotechnology | sc-32233 | 1:5,000 |
| Primary antibodies for immunofluorescence | MMP-12 – mouse monoclonal | Santa Cruz Biotechnology | sc-390863 | 1:100 |
| | GFAP – mouse monoclonal | Santa Cruz Biotechnology | sc-59992 | 1:100 |
| | Iba1 – mouse monoclonal | Thermo Fisher Scientific | MA5-27726 | 1:100 |
| | tPA – rabbit monoclonal | Novus Biologicals | NBP3-16356 | 1:100 |
| | PGRN – sheep polyclonal | R&D Systems | AF2557 | 1:6.7 |
| Secondary antibodies for immunoblotting | Goat anti-mouse IgG (H+L) – HRP | Novus Biologicals | NB7539 | 1:1,667 – 1:5,000 |
| | Goat anti-rabbit IgG (H+L) – HRP | Novus Biologicals | NB7160 | 1:1,667 – 1:5,000 |
| | Donkey anti-sheep IgG – HRP | R&D Systems | HAF016 | 1:1,667 – 1:5,000 |
| Secondary antibodies for immunofluorescence | Goat anti-mouse IgG (H+L) – Alexa Fluor 488 | Thermo Fisher Scientific | A-11001 | 1:200 |
| | Goat anti-rabbit IgG (H+L) – Alexa Fluor 594 | Thermo Fisher Scientific | A-11012 | 1:200 |
| | Donkey anti-sheep IgG (H+L) – Alexa Fluor 594 | Thermo Fisher Scientific | A-11016 | 1:200 |

MMP, matrix metalloproteinase; NA, not applicable; IFN, interferon; Iba, ionized calcium-binding adaptor molecule; MPO, myeloperoxidase; GFAP, glial fibrillary acidic protein; IL, interleukin; TNF, tumor necrosis factor; tPA, tissue-type plasminogen activator; PGRN, progranulin; GAPDH, glyceraldehyde-3-phosphate dehydrogenase; HRP, horseradish peroxidase.

**Supplementary Table 4.** Statistical tests and outcomes

| **Experimental groups** | **Statistical Test** | **Test outcomes with exact p values** |
|---|---|---|
| Figure 1A | | |
| Sham, Stroke (MMP-1) | Unpaired t test (two-tailed) - multiple comparisons | t ratio = 5.407<br>FDR-adjusted $p = 0.0004$ |
| Sham, Stroke (MMP-2) | Unpaired t test (two-tailed) - multiple comparisons | t ratio = 12.99<br>FDR-adjusted $p < 0.0001$ |
| Sham, Stroke (MMP-3) | Unpaired t test (two-tailed) - multiple comparisons | t ratio = 5.326<br>FDR-adjusted $p = 0.0004$ |
| Sham, Stroke (MMP-7) | Unpaired t test (two-tailed) - multiple comparisons | t ratio = 7.821<br>FDR-adjusted $p < 0.0001$ |
| Sham, Stroke (MMP-8) | Unpaired t test (two-tailed) - multiple comparisons | t ratio = 10.74<br>FDR-adjusted $p < 0.0001$ |
| Sham, Stroke (MMP-9) | Unpaired t test (two-tailed) - multiple comparisons | t ratio = 5.775<br>FDR-adjusted $p = 0.0003$ |
| Sham, Stroke (MMP-10) | Unpaired t test (two-tailed) - multiple comparisons | t ratio = 5.149<br>FDR-adjusted $p = 0.0005$ |
| Sham, Stroke (MMP-11) | Unpaired t test with Welch's correction (two-tailed) - multiple comparisons | t ratio = 10.46<br>FDR-adjusted $p < 0.0001$ |
| Sham, Stroke (MMP-12) | Unpaired t test (two-tailed) - multiple comparisons | t ratio = 12.15<br>FDR-adjusted $p < 0.0001$ |
| Sham, Stroke (MMP-13) | Unpaired t test (two-tailed) - multiple comparisons | t ratio = 7.596<br>FDR-adjusted $p < 0.0001$ |
| Sham, Stroke (MMP-14) | Unpaired t test (two-tailed) - multiple comparisons | t ratio = 14.3<br>FDR-adjusted $p < 0.0001$ |
| Sham, Stroke (MMP-15) | Unpaired t test (two-tailed) - multiple comparisons | t ratio = 5.597<br>FDR-adjusted $p = 0.0003$ |
| Sham, Stroke (MMP-16) | Unpaired t test (two-tailed) - multiple comparisons | t ratio = 2.850<br>FDR-adjusted $p = 0.0190$ |
| Sham, Stroke (MMP-17) | Unpaired t test (two-tailed) - multiple comparisons | t ratio = 2.381<br>FDR-adjusted $p = 0.0405$ |
| Sham, Stroke (MMP-19) | Unpaired t test (two-tailed) - multiple comparisons | t ratio = 15.02<br>FDR-adjusted $p < 0.0001$ |
| Sham, Stroke (MMP-20) | Unpaired t test (two-tailed) - multiple comparisons | t ratio = 12.28<br>FDR-adjusted $p < 0.0001$ |

| | | |
|---|---|---|
| Sham, Stroke (MMP-21) | Unpaired t test (two-tailed) - multiple comparisons | t ratio = 12.01<br>FDR-adjusted $p < 0.0001$ |
| Sham, Stroke (MMP-23) | Unpaired t test (two-tailed) - multiple comparisons | t ratio = 12.17<br>FDR-adjusted $p < 0.0001$ |
| Sham, Stroke (MMP-24) | Unpaired t test with Welch's correction (two-tailed) - multiple comparisons | t ratio = 3.677<br>FDR-adjusted $p = 0.0006$ |
| Sham, Stroke (MMP-25) | Mann-Whitney test (two-tailed) – multiple comparisons | Mann-Whitney U = 0.000<br>FDR-adjusted $p = 0.0001$ |
| Sham, Stroke (MMP-27) | Unpaired t test (two-tailed) - multiple comparisons | t ratio = 10.77<br>FDR-adjusted $p < 0.0001$ |
| Sham, Stroke (MMP-28) | Unpaired t test (two-tailed) - multiple comparisons | t ratio = 9.954<br>FDR-adjusted $p < 0.0001$ |
| Figure 1B | | |
| Stroke (MMP-12 vs. mNSS) | Spearman correlation test | r = 0.7047; $p = 0.013$ |
| Figure 1C | | |
| Sham, Stroke (MMP-12) | Two-way ANOVA | Interaction: F (1, 20) = 17.40; $p = 0.0005$<br>Brain Region: F (1, 20) = 16.36; $p = 0.0006$<br>Treatment: F (1, 20) = 25.24; $p < 0.0001$ |
| | Sidak's multiple comparisons test | Contralateral (Sham vs. Stroke): $p = 0.8003$<br>Ipsilateral (Sham vs. Stroke): $p < 0.0001$<br>Sham (Contralateral vs. Ipsilateral): $p = 0.9951$<br>Stroke (Contralateral vs. Ipsilateral): $p < 0.0001$ |
| Figure 1D | | |
| Sham, Stroke (MMP-12) | Unpaired t test with Welch's correction (two-tailed) | t = 14.98; $p < 0.0001$ |
| Figure 1E | | |
| Ctrl-sh, M12-sh (MMP-12) | Unpaired t test (two-tailed) | t = 6.493; $p = 0.0006$ |
| Figure 2A | | |
| Sham, Stroke, M12-sh | One-way ANOVA | F (2, 15) = 10.62; $p = 0.0013$ |
| | Tukey's multiple comparisons test | Sham vs. Stroke: $p = 0.0011$<br>Sham vs. M12-sh: $p = 0.2946$<br>Stroke vs. M12-sh: $p = 0.0240$ |
| Figure 2B | | |
| Stroke, M12-sh (GFAP) | Unpaired t test (two-tailed) - multiple comparisons | t ratio = 5.816<br>FDR-adjusted $p = 0.0007$ |

| | | |
|---|---|---|
| Stroke, M12-sh (Vimentin) | Unpaired t test with Welch's correction (two-tailed) - multiple comparisons | t ratio = 3.620<br>FDR-adjusted $p = 0.0160$ |
| Stroke, M12-sh (C3) | Unpaired t test with Welch's correction (two-tailed) - multiple comparisons | t ratio = 2.494<br>FDR-adjusted $p = 0.0488$ |
| Stroke, M12-sh (GBP2) | Unpaired t test (two-tailed) - multiple comparisons | t ratio = 4.910<br>FDR-adjusted $p = 0.0013$ |
| Figure 3B | | |
| Sham, Stroke, M12-sh | Brown-Forsythe ANOVA | F (2, 7.647) = 9.557; $p = 0.0083$ |
| | Dunnett's T3 multiple comparisons test | Sham vs. Stroke: $p = 0.0009$<br>Sham vs. M12-sh: $p = 0.0889$<br>Stroke vs. M12-sh: $p = 0.7678$ |
| Figure 3C | | |
| Ctrl-sh, M12-sh (IFNγ) | Unpaired t test (two-tailed) | t = 8.819; $p = 0.0001$ |
| Figure 3D | | |
| Stroke, M12-sh (CD68) | Unpaired t test (two-tailed) - multiple comparisons | t ratio = 5.152<br>FDR-adjusted $p = 0.0023$ |
| Stroke, M12-sh (TLR2) | Unpaired t test (two-tailed) - multiple comparisons | t ratio = 3.229<br>FDR-adjusted $p = 0.0119$ |
| Stroke, M12-sh (TLR4) | Unpaired t test with Welch's correction (two-tailed) - multiple comparisons | t ratio = 3.355<br>FDR-adjusted $p = 0.0039$ |
| Stroke, M12-sh (IL-1β) | Unpaired t test (two-tailed) - multiple comparisons | t ratio = 3.251<br>FDR-adjusted $p = 0.0119$ |
| Stroke, M12-sh (IL-6) | Unpaired t test with Welch's correction (two-tailed) - multiple comparisons | t ratio = 2.370<br>FDR-adjusted $p = 0.0120$ |
| Figure 3E | | |
| Ctrl-sh, M12-sh (IL-1β) | Unpaired t test (two-tailed) | t = 9.510; $p < 0.0001$ |
| Ctrl-sh, M12-sh (IL-6) | Unpaired t test (two-tailed) | t = 6.056; $p = 0.0009$ |
| Ctrl-sh, M12-sh (TNFα) | Unpaired t test (two-tailed) | t = 7.177; $p = 0.0004$ |
| Figure 4A | | |
| Sham, Stroke (CCR2) | Unpaired t test (two-tailed) – multiple comparisons | t ratio = 11.45<br>FDR-adjusted $p < 0.0001$ |
| Sham, Stroke (CXCR2) | Unpaired t test (two-tailed) – multiple comparisons | t ratio = 10.29<br>FDR-adjusted $p < 0.0001$ |

| | | |
|---|---|---|
| Sham, Stroke (ELANE) | Unpaired t test (two-tailed) – multiple comparisons | t ratio = 6.768<br>FDR-adjusted $p < 0.0001$ |
| Sham, Stroke (MPO) | Unpaired t test (two-tailed) – multiple comparisons | t ratio = 4.623<br>FDR-adjusted $p = 0.0010$ |
| Figure 4B | | |
| Stroke (MMP-12 vs. CCR2) | Pearson correlation test | r = 0.8631; $p = 0.0003$ |
| Stroke (MMP-12 vs. CXCR2) | Pearson correlation test | r = 0.7244; $p = 0.0077$ |
| Stroke (MMP-12 vs. MPO) | Spearman correlation test | r = 0.7692; $p = 0.0049$ |
| Figure 4C | | |
| Sham, Stroke (MPO) | Unpaired t test (two-tailed) | t = 22.64; $p < 0.0001$ |
| Figure 4D | | |
| Ctrl-sh, M12-sh (MPO) | Unpaired t test | t = 6.319; $p = 0.0007$ |
| Figure 5 | | |
| Stroke, M12-sh (MMP-2) | Unpaired t test with Welch's correction (two-tailed) - multiple comparisons | t ratio = 4.774<br>FDR-adjusted $p = 0.0033$ |
| Stroke, M12-sh (MMP-3) | Unpaired t test (two-tailed) - multiple comparisons | t ratio = 2.478<br>FDR-adjusted $p = 0.0129$ |
| Stroke, M12-sh (MMP-7) | Unpaired t test (two-tailed) - multiple comparisons | t ratio = 1.702<br>FDR-adjusted $p = 0.0396$ |
| Stroke, M12-sh (MMP-8) | Unpaired t test (two-tailed) - multiple comparisons | t ratio = 1.757<br>FDR-adjusted $p = 0.0383$ |
| Stroke, M12-sh (MMP-9) | Unpaired t test (two-tailed) - multiple comparisons | t ratio = 2.969<br>FDR-adjusted $p = 0.0078$ |
| Stroke, M12-sh (MMP-11) | Unpaired t test with Welch's correction (two-tailed) - multiple comparisons | t ratio = 5.641<br>FDR-adjusted $p = 0.0027$ |
| Stroke, M12-sh (MMP-14) | Unpaired t test (two-tailed) - multiple comparisons | t ratio = 7.839<br>FDR-adjusted $p < 0.0001$ |
| Stroke, M12-sh (MMP-15) | Unpaired t test (two-tailed) - multiple comparisons | t ratio = 4.762<br>FDR-adjusted $p = 0.0008$ |
| Stroke, M12-sh (MMP-19) | Unpaired t test (two-tailed) - multiple comparisons | t ratio = 7.005<br>FDR-adjusted $p = 0.0001$ |
| Stroke, M12-sh (MMP-20) | Unpaired t test (two-tailed) - multiple comparisons | t ratio = 1.342<br>FDR-adjusted $p = 0.0629$ |

| | | |
|---|---|---|
| Stroke, M12-sh (MMP-21) | Unpaired t test (two-tailed) - multiple comparisons | t ratio = 2.939<br>FDR-adjusted $p = 0.0078$ |
| Stroke, M12-sh (MMP-23) | Unpaired t test (two-tailed) - multiple comparisons | t ratio = 6.512<br>FDR-adjusted $p = 0.0001$ |
| Stroke, M12-sh (MMP-25) | Mann-Whitney test (two-tailed) – multiple comparisons | Mann-Whitney U = 1.000<br>FDR-adjusted $p = 0.0040$ |
| Stroke, M12-sh (MMP-27) | Unpaired t test (two-tailed) - multiple comparisons | t ratio = 2.695<br>FDR-adjusted $p = 0.0106$ |
| Stroke, M12-sh (MMP-28) | Mann-Whitney test (two-tailed) – multiple comparisons | Mann-Whitney U = 3.000<br>FDR-adjusted $p = 0.0086$ |
| Figure 6C | | |
| Ctrl-sh, M12-sh (tPA) | Unpaired t test (two-tailed) | t = 4.237; $p = 0.0055$ |
| Figure 7A | | |
| Sham, Stroke (PGRN) | Unpaired t test (two-tailed) | t = 32.81; $p < 0.0001$ |
| Figure 7B | | |
| Sham, Stroke (PGRN-80 kDa) | Unpaired t test | t = 0.1288; $p = 0.9007$ |
| Sham, Stroke (PGRN-68 kDa) | Unpaired t test with Welch's correction (two-tailed) | t = 3.767; $p = 0.0193$ |
| Figure 7C | | |
| Stroke (MMP-12 vs. PGRN) | Pearson correlation test | r = 0.6100; $p = 0.0352$ |
| Figure 7E | | |
| Stroke, M12-sh (PGRN) | Unpaired t test with Welch's correction (two-tailed) | t = 5.000; $p = 0.0027$ |
| Figure 7F | | |
| Ctrl-sh, M12-sh (PGRN-80 kDa) | Unpaired t test | t = 3.128; $p = 0.0204$ |
| Ctrl-sh, M12-sh (PGRN-68 kDa) | Unpaired t test | t = 2.668; $p = 0.0371$ |
| Supplementary Figure 1 | | |
| Sham, Stroke (GFAP) | Unpaired t test (two-tailed) - multiple comparisons | t ratio = 15.95<br>FDR-adjusted $p < 0.0001$ |
| Sham, Stroke (Vimentin) | Unpaired t test (two-tailed) - multiple comparisons | t ratio = 31.17<br>FDR-adjusted $p < 0.0001$ |

| | | |
|---|---|---|
| Sham, Stroke (C3) | Mann-Whitney test (two-tailed) – multiple comparisons | Mann-Whitney U = 0.000<br>FDR-adjusted $p = 0.0023$ |
| Sham, Stroke (GBP2) | Unpaired t test (two-tailed) - multiple comparisons | t ratio = 24.36<br>FDR-adjusted $p < 0.0001$ |
| Sham, Stroke (FKBP5) | Mann-Whitney test (two-tailed) – multiple comparisons | Mann-Whitney U = 0.000<br>FDR-adjusted $p = 0.0023$ |
| Supplementary Figure 2 | | |
| Stroke (MMP-12 vs. GFAP) | Spearman correlation test | r = 0.6713; $p = 0.0202$ |
| Stroke (MMP-12 vs. Vimentin) | Pearson correlation test | r = 0.6431; $p = 0.0241$ |
| Stroke (MMP-12 vs. C3) | Pearson correlation test | r = 0.6230; $p = 0.0304$ |
| Stroke (MMP-12 vs. GBP2) | Pearson correlation test | r = 0.7972; $p = 0.0029$ |
| Supplementary Figure 3 | | |
| Sham, Stroke (IFNγ) | Unpaired t test with Welch's correction (two-tailed) | t = 12.91; $p = 0.0001$ |
| Supplementary Figure 4 | | |
| Sham, Stroke (CD68) | Unpaired t test (two-tailed) - multiple comparisons | t ratio = 24.27<br>FDR-adjusted $p < 0.0001$ |
| Sham, Stroke (CD16) | Unpaired t test (two-tailed) - multiple comparisons | t ratio = 25.54<br>FDR-adjusted $p < 0.0001$ |
| Sham, Stroke (TLR2) | Unpaired t test (two-tailed) - multiple comparisons | t ratio = 15.33<br>FDR-adjusted $p < 0.0001$ |
| Sham, Stroke (TLR4) | Unpaired t test (two-tailed) - multiple comparisons | t ratio = 11.05<br>FDR-adjusted $p < 0.0001$ |
| Sham, Stroke (iNOS) | Unpaired t test (two-tailed) - multiple comparisons | t ratio = 9.805<br>FDR-adjusted $p < 0.0001$ |
| Sham, Stroke (CCL2) | Unpaired t test (two-tailed) - multiple comparisons | t ratio = 22.26<br>FDR-adjusted $p < 0.0001$ |
| Sham, Stroke (CXCL1) | Unpaired t test with Welch's correction (two-tailed) - multiple comparisons | t ratio = 8.911<br>FDR-adjusted $p = 0.0001$ |
| Sham, Stroke (IL-1β) | Mann-Whitney test (two-tailed) – multiple comparisons | Mann-Whitney U = 5.000<br>FDR-adjusted $p = 0.0432$ |

| | | |
|---|---|---|
| Sham, Stroke (IL-6) | Unpaired t test (two-tailed) - multiple comparisons | t ratio = 8.369<br>FDR-adjusted $p < 0.0001$ |
| Sham, Stroke (TNFα) | Unpaired t test (two-tailed) - multiple comparisons | t ratio = 4.971<br>FDR-adjusted $p = 0.0007$ |
| Supplementary Figure 5 | | |
| Sham, Stroke (IL-1β) | Unpaired t test with Welch's correction (two-tailed) | t = 10.78; $p = 0.0002$ |
| Sham, Stroke (IL-6) | Unpaired t test with Welch's correction (two-tailed) | t = 10.66; $p = 0.0004$ |
| Sham, Stroke (TNFα) | Unpaired t test with Welch's correction (two-tailed) | t = 10.13; $p = 0.0004$ |
| Supplementary Figure 6 | | |
| Stroke (MMP-12 vs. CCL2) | Spearman correlation test | r = 0.7552; $p = 0.0062$ |
| Stroke (MMP-12 vs. CXCL1) | Spearman correlation test | r = 0.8182; $p = 0.0019$ |
| Stroke (MMP-12 vs. IL-6) | Spearman correlation test | r = 0.7692; $p = 0.0049$ |
| Supplementary Figure 7 | | |
| Sham, Stroke (CD163) | Mann-Whitney test (two-tailed) – multiple comparisons | Mann-Whitney U = 1.000<br>FDR-adjusted $p = 0.0045$ |
| Sham, Stroke (CD206) | Unpaired t test (two-tailed) - multiple comparisons | t ratio = 11.04<br>FDR-adjusted $p < 0.0001$ |
| Sham, Stroke (Arg1) | Unpaired t test with Welch's correction (two-tailed) - multiple comparisons | t ratio = 7.847<br>FDR-adjusted $p = 0.0005$ |
| Sham, Stroke (IL-10) | Unpaired t test (two-tailed) - multiple comparisons | t ratio = 8.125<br>FDR-adjusted $p < 0.0001$ |
| Sham, Stroke (TGFβ) | Unpaired t test (two-tailed) - multiple comparisons | t ratio = 21.34<br>FDR-adjusted $p < 0.0001$ |
| Supplementary Figure 8 | | |
| Stroke (MMP-12 vs. CD163) | Pearson correlation test | r = 0.7468; $p = 0.0053$ |
| Stroke (MMP-12 vs. CD206) | Pearson correlation test | r = 0.7592; $p = 0.0042$ |
| Stroke (MMP-12 vs. Arg1) | Pearson correlation test | r = 0.7645; $p = 0.0038$ |

| | | |
|---|---|---|
| Stroke (MMP-12 vs. IL-10) | Spearman correlation test | r = 0.6434; *p* = 0.0278 |
| Stroke (MMP-12 vs. TGFβ) | Pearson correlation test | r = 0.7182; *p* = 0.0085 |
| Supplementary Figure 9 | | |
| Stroke, M12-sh (CD163) | Mann-Whitney test (two-tailed) – multiple comparisons | Mann-Whitney U = 5.000<br>FDR-adjusted *p* = 0.0205 |
| Stroke, M12-sh (CD206) | Unpaired t test (two-tailed) - multiple comparisons | t ratio = 1.959<br>FDR-adjusted *p* = 0.0401 |
| Stroke, M12-sh (Arg1) | Unpaired t test (two-tailed) - multiple comparisons | t ratio = 3.138<br>FDR-adjusted *p* = 0.0074 |
| Stroke, M12-sh (IL-10) | Unpaired t test (two-tailed) - multiple comparisons | t ratio = 3.341<br>FDR-adjusted *p* = 0.0074 |
| Stroke, M12-sh (TGFβ) | Unpaired t test (two-tailed) - multiple comparisons | t ratio = 7.868<br>FDR-adjusted *p* < 0.0001 |

MMP, matrix metalloproteinase; mNSS, modified Neurological Severity Score; FDR, False Discovery rate; GFAP, glial fibrillary acidic protein; C3, complement component 3; GBP2, guanylate-binding protein 2; FKBP5, FK506 binding protein 5; IFN, interferon; TLR, toll-like receptor; iNOS, inducible nitric oxide synthase; CCL2, C-C motif chemokine ligand 2; CXCL1, C-X-C motif chemokine ligand 1; IL, interleukin; TNF, tumor necrosis factor; CCR2, C-C motif chemokine receptor 2; CXCR2, C-X-C Motif Chemokine Receptor 2; ELANE, elastase, neutrophil expressed; MPO, myeloperoxidase; tPA, tissue-type plasminogen activator; PGRN, progranulin; Arg1, arginase 1; TGFβ, Transforming growth factor β.
